# Substrate-engaged type III secretion system structures reveal gating mechanism for unfolded protein translocation

**DOI:** 10.1101/2020.12.17.423328

**Authors:** Sean Miletic, Dirk Fahrenkamp, Nikolaus Goessweiner-Mohr, Jiri Wald, Maurice Pantel, Oliver Vesper, Vadim Kotov, Thomas C. Marlovits

## Abstract

Many bacterial pathogens strictly rely on the activity of type III secretion systems (T3SSs) to secrete and translocate effector proteins in order to establish infection. The central component of T3SSs is the needle complex, a supramolecular machine which assembles a continuous conduit crossing the bacterial envelope and the host cell membrane to allow bacterial effectors to gain entry into the host cell cytoplasm to modulate signal transduction processes. Disruption of this process impairs pathogenicity, providing an avenue for antimicrobial design. However, the molecular principles underlying T3 secretion remain elusive. Here, we report the first structure of an active *Salmonella enterica* sv. Typhimurium needle complex engaged with the late effector protein SptP in two functional states, revealing the complete 800Å-long secretion conduit and unravelling the critical role of the export apparatus (EA) subcomplex in T3 secretion. Unfolded substrates enter the EA through a hydrophilic constriction formed by SpaQ proteins, which enables side chain-independent transport, explaining heterogeneity and structural disorder of signal sequences in T3SS effector proteins. Above, a methionine gasket formed by SpaP proteins functions as a gate that dilates to accommodate substrates but prevents leaky pore formation to maintain the physical boundaries of compartments separated by a biological membrane. Following gate penetration, a moveable SpaR loop first folds up to then act akin to a linear ratchet to steer substrates through the needle complex. Together, these findings establish the molecular basis for substrate translocation through T3SSs, improving our understanding of bacterial pathogenicity and motility of flagellated bacteria, and paves the way for the development of novel concepts combating bacterial infections.

## Main Text

Many important human pathogens including *Salmonella, Shigella, Yersinia*, and enteropathogenic *Escherichia coli* (EPEC) employ a conserved type III secretion system (T3SS) also termed injectisome to deliver a pleiotropic arsenal of proteins into target eukaryotic cells^1^. These proteins modulate host cell signal transduction processes to establish a biological niche within the host, making T3SSs crucial virulence determinants^2^. Yet, the precise mechanisms that allow these secretion systems to facilitate unfolded protein transport across the bacterial envelope and into the host cell while maintaining bacterial integrity remain poorly understood. Therefore, visualizing the translocation process at the molecular level is essential for our understanding of host-pathogen biology and the development of novel therapies targeting bacterial infection.

The T3SS is a large molecular machine, over 3.6 megadaltons in mass, spanning across the inner and outer bacterial membranes with an extracellular filamentous appendage extending out to target host cells. Chaperones present effector proteins in a non-globular, secretion-competent state to a cytoplasmic sorting platform complex, which sorts and loads effectors into the export apparatus (EA) subcomplex located inside the membrane-bound basal body^3,4,5,6^. Extending from the EA is a long, helical needle filament, capped by a tip complex that contacts the host cell membrane via assembly of a translocon pore^7,8,9^. The basal body and the needle filament, collectively termed the needle complex, function as a continuous conduit for effector protein translocation from the prokaryotic to the host cell cytoplasm^10,11^.

Accumulating structural information has revealed a shared common architecture between virulent and flagellar T3SSs, especially in the EA^6,12–15^. However, in all structures known to date, the proposed translocation channel through the EA is sealed by a gasket with an above loop, making comprehension of substrate transport through the needle complex difficult. Furthermore, it remains unclear how the EA achieves selective effector protein transport given the multitude of proteins that are present in the bacterial cytoplasm.

Visualizing actively-secreting injectisomes is however difficult due to the rapid dynamics of protein transport. A pool of protein substrates is translocated through the T3SS in a hierarchical order upon host cell contact, although injectisomes can be artificially induced to secrete proteins *in vitro^4,16^.* It is unclear what proportion of these injectisomes actively secrete proteins as in *Salmonella,* induced cells can contain tens of needle complexes^10^. Furthermore, translocation is rapid, estimated at a rate of 7-60 molecules per second^17^. With this speed and temporal variability, isolated needle complexes likely lack protein substrates, or they dissociate during purification procedures. To overcome these hurdles, effector proteins can be artificially trapped in needle complexes by fusion to C-terminal tags resistant to unfolding^18,19^. We previously showed that the *Salmonella* late-translocated effector protein SptP fused to a GFP tag can be visualized as a subtracted density in the needle complex, confirming that the filament functions as the conduit for effector proteins^18^. However, direct visualization of a substrate throughout the complete secretion conduit has remained challenging, leaving the questions as to how and where the EA would eventually open to allow passage of effector proteins, while maintaining the integrity and composition of compartments separated by a biological membrane, unresolved.

## Results

To obtain molecular snapshots of a T3SS engaged with a substrate, we applied cryo-EM to purified injectisome complexes which had been enriched for trapped SptP3x-GFP by immunoprecipitation (Supplementary Figs. 1 and 2). Single particle reconstruction provided us with a non-symmetrized density map of the substrate-trapped needle complex in two active functional states, ranging from 2.4 to 4.5Å in resolution (Fig. 1, Supplementary Figs. 3-5, Supplementary Table 1), and resolving a substrate density traversing through the complete secretion path from the cytoplasmic face of the needle complex to the extracellular filament (Fig. 1a,b). The SptP density reveals that the effector protein adopts a non-globular fold during transport through the needle complex (Fig. 1). However, the positional and conformational flexibility of the substrate, propagated throughout the entire translocation path, impeded our efforts to assign specific residues. As a consequence, we modeled the SptP3x-GFP substrate as a polyalanine sequence (Supplementary Fig. 6).

**Fig. 1.**
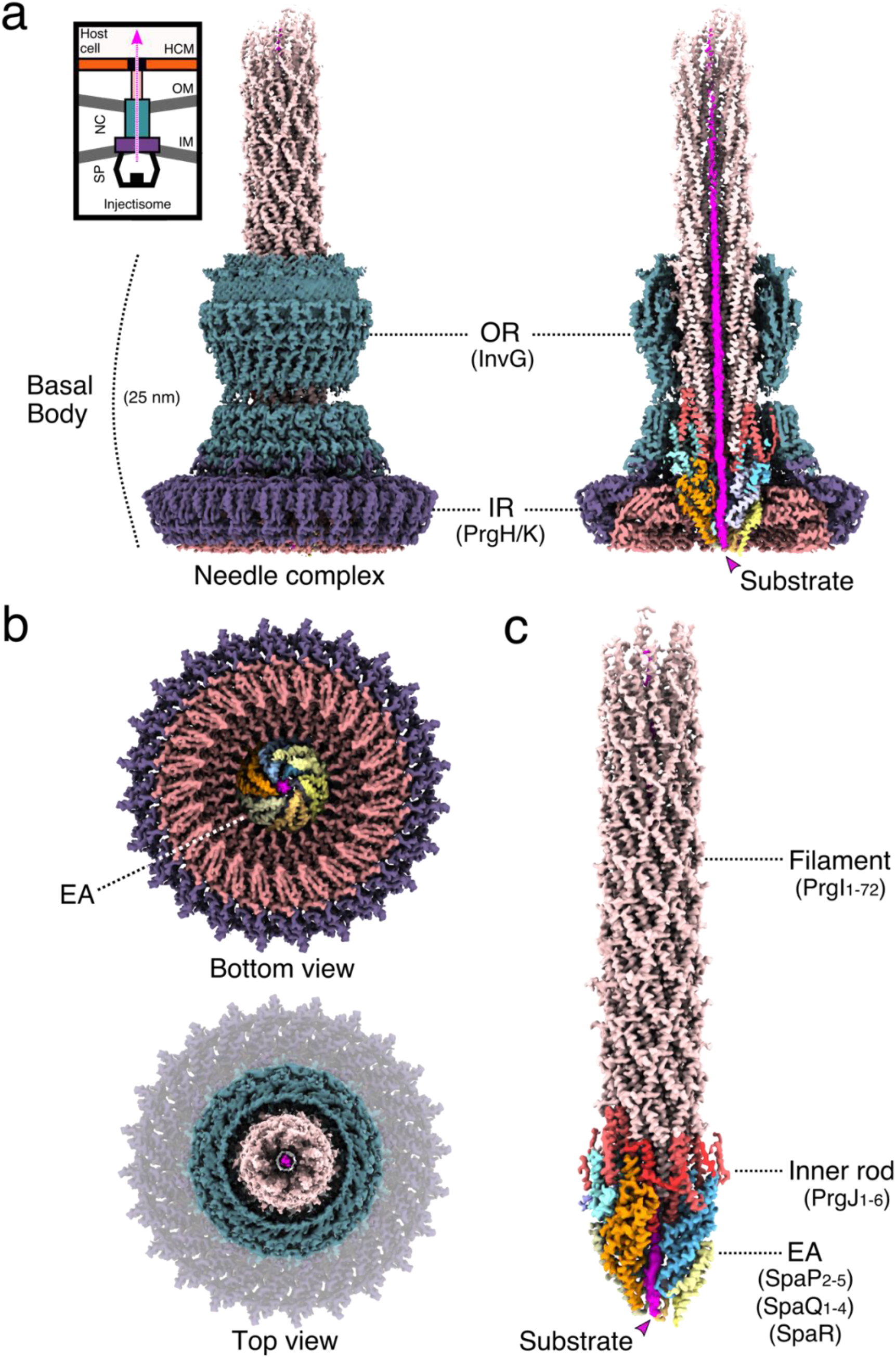
CryoEM map of the *S. enterica* sv. Typhimurium needle complex engaged with the effector protein substrate SptP3x-GFP. **a**, The non-symmetrized cryoEM (C1) reconstruction of the substrate-engaged needle complex (left) with a vertical cross section through the center (right) revealing the substrate shown in magenta throughout the translocation channel. Upper left, is a cartoon schematic of the needle complex injectisome in the bacterial inner and outer membranes (IM/OM) and in contact with a host cell membrane (HCM). NC: needle complex, SP: sorting platform, OR: outer rings, IR: inner rings. PrgH_1-24_: dark purple, PrgK_1-24_: amaranth, InvG_1-15/16_: dark green, SpaQ_1-4_: yellow colors, SpaP_1-5_: blue colors, SpaR: orange, PrgJ_1-6_: red, PrgI_1-72_: salmon. **b**, Top and bottom views of the C1 map showing the export apparatus (EA) and substrate. **c**, CryoEM map of the filament, inner rod and the export apparatus components. SpaP1 has been removed to aid visualization of the substrate.

Inside the needle complex, the substrate travels through a secretion conduit built by the EA, a decameric subcomplex made up of three proteins, SpaP (5x), SpaQ (4x), and SpaR (1x), the inner rod composed of PrgJ1-6, and the PrgI-containing filament (Figs. 1c and 2a). Together, these proteins form three discrete building blocks that are embedded within three oligomeric protein rings formed by InvG, PrgH and PrgK, a scaffold spanning the two bacterial membranes and the periplasm (Fig. 1a,b and Supplementary Fig. 7).

**Fig. 2.**
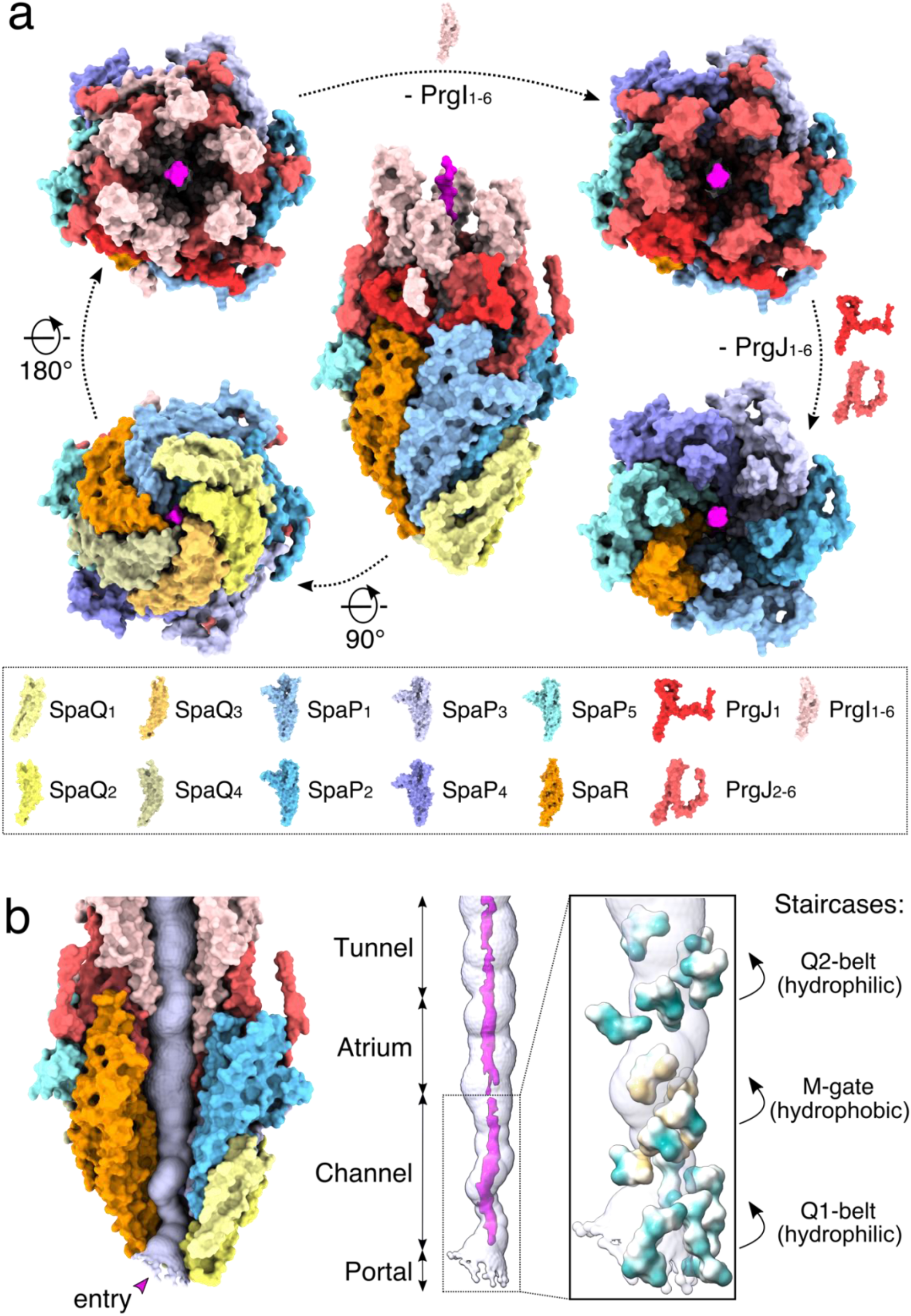
The export apparatus (EA) forms a translocation channel for substrates. **a**, Modular assembly of the substrate-trapped EA, inner rod (PrgJ) and the first six PrgI subunits of the helical filament. Individual protein components are shown in the dashed box below. SptP3x-GFP is shown in magenta. **b**, Left: view of the EA with SpaP1, PrgJ1-2 and PrgI1 removed and the substrate translocation path displayed as a white surface. Right: four discrete sections (portal, channel, atrium and tunnel) of the translocation path are shown with the EM density corresponding to the substrate (threshold: 0.015). Right box: magnification highlighting surfaces of residues forming hydrophilic and hydrophobic staircases encircling the portal and channel. Green: hydrophilic; white: neutral; gold: hydrophobic.

The EA can be further separated into three discrete sections which form a three-point pseudo-helical interface with the substrate that together consists of two hydrophilic constrictions containing conserved glutamine residues (hereafter referred to as Q1- and Q2-belt) that sandwich a hydrophobic methionine gasket (hereafter referred to as M-gate). The substrate enters the EA with its N-terminus through the portal containing the Q1-belt, continues through the M-gate and Q2-belt defining the EA channel, before reaching the atrium chamber of the inner rod and finally the filament tunnel (Fig. 2b).

To be able to investigate the structural changes underlying substrate transport, we also determined the structure of an apo-state complex. Focused refinement without any symmetry enforcement provided us with a reconstruction yielding an average resolution of ~3.3Å and resolving the entire EA, the inner rod and parts of the filament (Supplementary Fig. 8). The model that we built is in very good agreement with a published apo-state structure of the same complex (6pep) and structurally-related complexes (6r6b, 6r69, 6s3r, 6s3l, 6s3s), together showing (i) closed EAs share a conserved architecture with a defined 5:4:1 (SpaP:Q:R) stoichiometry and (ii) suggesting that a conserved mechanism orchestrates substrate translocation through these secretion systems (Supplementary Figs. 9 and 10)^6,12,13^. Intriguingly, our apo- and translocation-state EAs superimpose with a very low root mean square deviation (rmsd) of 0.49Å (SpaP:Q:R), revealing that in fact only subtle conformational changes are needed to facilitate substrate transport through the channel of the needle complex (Supplementary Fig. 11).

The substrate first engages the needle complex structure through the EA core complex portal formed by four SpaQ proteins, confirming our previous results that the central opening localizing to the cytoplasmic tip of the EA serves as the substrate entry site^18^. Four SpaQ loops connecting α-helices α1 and α2 and a lone SpaR Gln208 together form the Q1-belt in the cytoplasmic tip of the EA core complex (Fig. 3a). SpaS, which binds to the EA complex and simultaneously wraps around all four SpaQ loops in the homologous and recombinantly produced EA structure from *Vibrio mimicus* (6s3l), dissociates from our substrate-engaged needle complex during purification, which could explain why SpaQ_1_, and to a lesser extent SpaQ_2_, adopt a more open conformation (rmsd: 1.27Å; Supplementary Fig. 12)^13^. SpaQ homologues of many important human pathogens share a conserved Gln-X-Gln-X-Gln motif within the aforementioned loop, which effectively renders the environment in the Q1-belt hydrophilic (Supplementary Fig. 13). The high sequence conservation strongly suggests that the Q1-belt plays an essential role for substrate translocation through bacterial T3SSs.

**Fig. 3.**
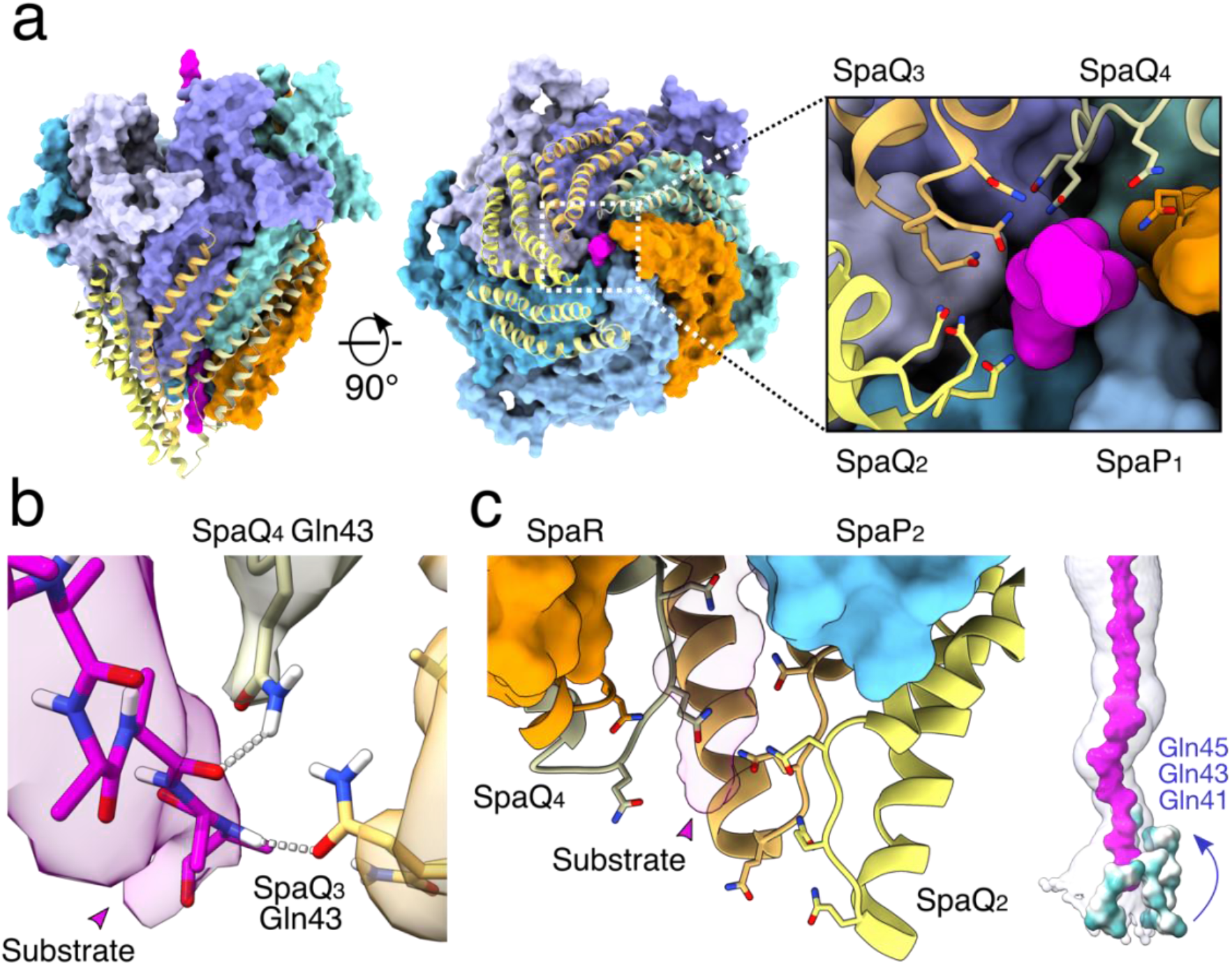
The Q1-belt portal of the EA facilitates substrate loading into the needle complex. **a**, Surface representation of the substrate-engaged EA, with SpaQ_1-4_ shown as yellow ribbon diagrams. Right box: a magnified view of the conserved SpaQ glutamine residues Gln41, Gln43 and Gln45 involved in substrate engagement, depicted in stick representation. **b**, Hydrogen-bond formation between the substrate backbone and SpaQ_3_ Gln43 and SpaQ_4_ Gln43. The surfaces represent EM density (threshold: 0.014). **c**, Side view of the SpaQ Q1-belt displayed as ribbon diagrams encircling the SptP3x-GFP substrate with the side chains of Gln41/43/45 shown in stick representation. Far right: Surface representations of the Q1 belt residues Gln41/43/45 colored by hydrophobicity. Green: hydrophilic; white: neutral; gold: hydrophobic.

Based on the appearance of its density, the substrate is largely unfolded in the Q1-belt area of the translocation path, indicating sequences corresponding to loops and β-sheets in natively folded SptP have mostly been trapped in the EA portal (Figs. 2b and 3b, Supplementary Fig. 6). Notably, little structural information is available for the very N-termini of T3SS effector proteins, especially when complexed with their cognate chaperones, supporting prediction models in which these sequences are typically intrinsically disordered^20,21^. Therefore, it is conceivable that our model provides mechanistic insights into how substrate proteins are loaded into the needle complex. In total, the Q1-belt is shaped by thirteen glutamines (3x in each of the four SpaQs, 1x in SpaR) that localize within close proximity to the trapped SptP3x-GFP substrate. Out of these, Gln43 of SpaQ_3_ and SpaQ_4_ establish hydrogen-bond interactions with the substrate backbone carbonyl oxygens and amine hydrogens (Fig. 3b). Due to the spiral staircase arrangement, the SpaQ/R glutamines provide complementary interaction interfaces over a length of ~20Å, which, upon successive binding to the substrate, cause an increase in avidity that stabilizes the substrate in the EA portal below the M-gate (Figs. 2 and 3b,c). Together, our structural data reveals that loading of effector proteins into the needle complex can be accomplished in a side chain-independent fashion and hence provides a rationale to explain the structural disorder and plasticity of N-terminal signal sequences observed in bacterial T3SS effector proteins.

Following engagement in the hydrophilic Q1-belt, the substrate asymmetrically twists up through the ~5.5Å-wide M-gate (Fig. 4a,b). The loops between the fifth and sixth alpha helix of each of the five SpaP proteins contain three conserved methionine residues, which, together with conserved SpaR Phe212, form a hydrophobic constriction seen in substrate-free structures^15,6,12,13^ (Fig. 4a,b and Supplementary Figs. 14-16). Similar to the Q1-belt, the pseudo helical structure of the EA causes these methionines to form a ~20Å-long, spiral staircase-like gate that shifts to accommodate the substrate passing through (Figs. 2 and 4a,b).

**Fig. 4.**
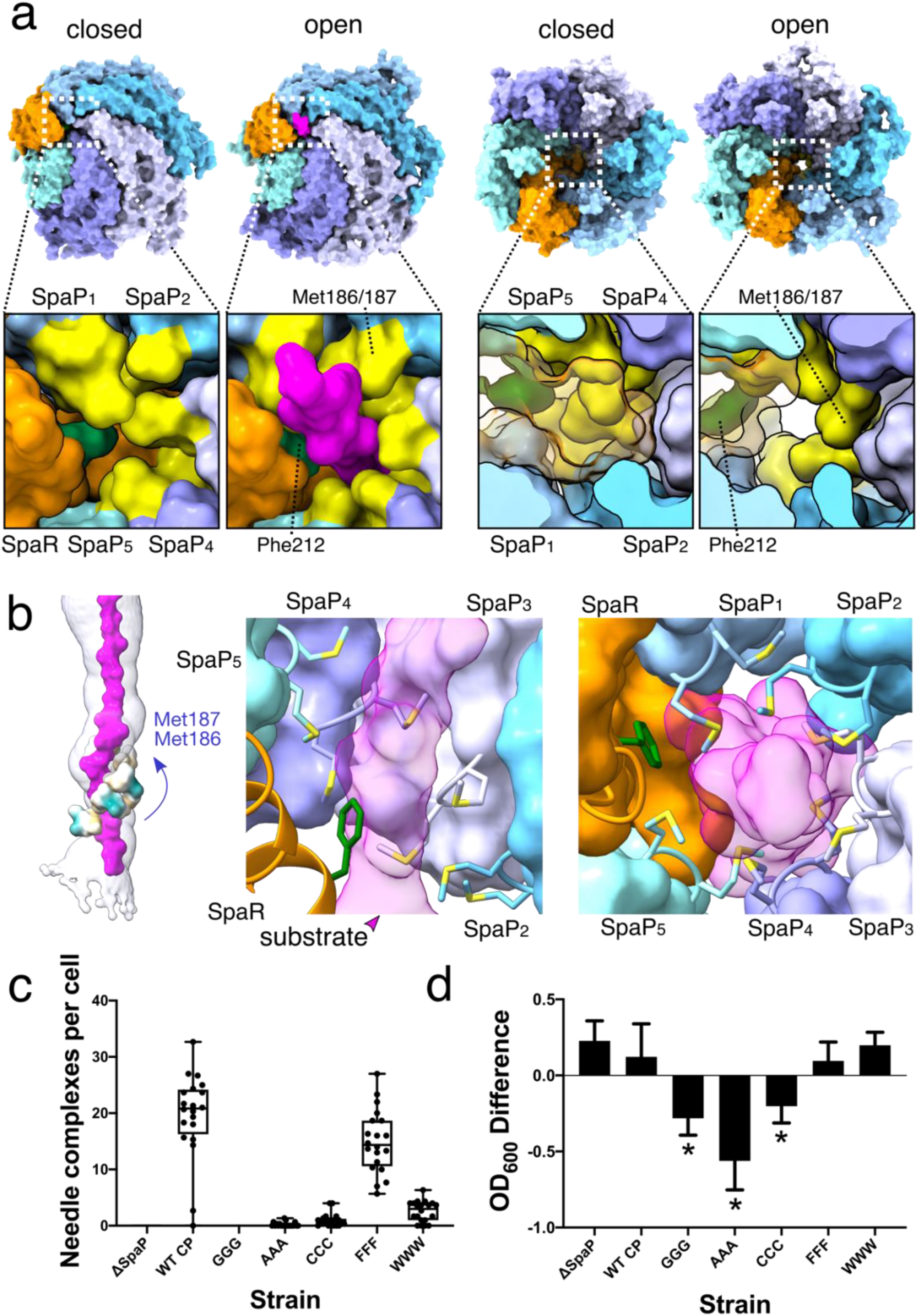
Opening of the M-gate in the EA facilitates substrate translocation and is crucial for needle complex assembly and cellular fitness. **a**, Bottom and top views of the EA, with M-gate shown in the lower panels, depicted in surface representation in a closed or open (substrate engaged) state. Methionine residues 186/7 are displayed in yellow and Phe212 in green. SpaR is displayed transparent in the top views. **b**, Left: surface representations of the M-gate residues Met186/7 colored by hydrophobicity. Middle and right: side and bottom views of the M-gate staircase with Met186/7 and Phe212 depicted in stick representation. **c**, Quantification of needle complexes (n=20 cells) in *Salmonella* SpaP knockout strains complemented with SpaP WT (WT CP) or with mutants targeting the conserved 185-Met-Met-Met-187 motif of the M-gate (GGG-WWW). Boxplot whiskers show min and max counts. Individual counts are represented by dots. Needle complexes were counted by three individuals and averaged. **d**, Optical densities of the *Salmonella* strains in (**c**). Plotted is the mean difference in OD_600_ between cultures grown for 6 hrs under T3SS-inducing and non-inducing conditions. Error bars represent SD and asterisks represent a significant difference compared to WT CP (GGG *P* < 0.0001, AAA *P* < 0.0001, CCC *P* = 0.002). A one-way ANOVA and a Dunnett’s test were used to assess statistical significance between the WT control (WT CP) and M-gate substitution strains.

Notably, the SpaP pentamer (SpaP1-5) in our substrate-engaged structure superimposes well with its apo-state counterpart (rmsd: 0.45Å), which reinforces the concept that transport of substrate proteins across the bacterial envelope does not involve large conformational changes but is facilitated by subtle rearrangements that, within the M-gate, are mostly limited to the side chains of the three methionines (Fig. 4a-b and Supplementary Fig. 11). Similar to the Q1-belt area, the density corresponding to the trapped substrate fits best to an unfolded polypeptide (Fig. 2b and Supplementary Fig. 6), which, together with the size constraints imposed by the M-gate (~5.5Å), supports our observation that non-folded SptP sequences have been trapped in the EA.

To functionally characterize the M-gate based on our structural data, we reconstituted a *Salmonella* SpaP knockout strain (SpaP^KO^) with exogenous SpaP carrying mutations targeting the conserved methionine motif. Negative stain EM revealed that substitution of the motif with the aliphatic amino acids glycine and alanine (SpaP^GGG^, SpaP^AAA^), as well as cysteine (SpaPCCC), resulted in markedly reduced numbers of needle complexes under injectisome-inducing conditions, demonstrating that needle complex assembly is impaired in these strains (Fig. 4c and Supplementary Fig. 17). Because proteins forming the inner rod (PrgJ) and the filament (PrgI) are transported through the T3SS as a natural part of the injectisome assembly process, our data provides evidence that at least the alanine and cysteine mutant strains retain some ability to actively transport proteins through their mutated EAs and hence the methionine motif alone is not strictly required for substrate translocation^22^. However, all three strains display impaired growth kinetics, suggesting that the assembly of EAs whose M-gates are composed of residues with side chains smaller than methionine creates pores in the inner bacterial membrane that likely short circuit the membrane potential and cause reduced pathogen fitness (Fig. 4d and Supplementary Figs. 18 and 19). Consistent with this hypothesis, deletion of a single methionine has been shown to be sufficient to increase membrane conductance in the flagellar homologue FliP^23^.

To further corroborate this hypothesis, we introduced either tryptophans (SpaP^WWW^) or phenylalanines (SpaP^FFF^) into SpaP to mimic the hydrophobic properties of methionine. Evidently, substitution of the methionine motif with these amino acids did not affect bacterial growth kinetics (Fig. 4d and Supplementary Fig. 18). However, substitution with phenylalanine but not tryptophan restored needle complex assembly to almost wild-type levels, indicating that the bulky hydrophobic side chain of tryptophan effectively prevents leaky pore formation but, compared to methionine and phenylalanine, appears to be too inflexible to efficiently facilitate substrate translocation (Fig. 4c, Supplementary Fig. 17).

Based on our findings, we reasoned that substrates penetrating the methionine network of the EA cause an opening of what appears to be a hydrophobic gate just large enough to accommodate the unfolded substrate chain. By intimately engaging the translocating substrate, the M-gate effectively acts as a tight seal to facilitate transport but maintain the physical boundaries between the pathogen’s cytoplasm and (i) its periplasm during needle complex assembly, (ii) the outside environment (prior to infection) and (iii) the host cell cytoplasm (during infection).

In the apo-state, a unique loop of residues 106-123 of SpaR extends horizontally out on top of the M-gate (Fig. 5a). Consistent with the recombinant export apparatus from *Shigella flexneri* injectisomes, SpaR residues Leu110 and Ile114 interface with the methionines of the M-gate below, creating what has been termed a ‘plug’ in the structurally-related flagellar system (Supplementary Fig. 20)^12,15^. In the map of our substrate-engaged complex, the SpaR loop or ‘lid’ adopts two distinct conformational states positioned vertically along the translocation path (Fig. 5a and Supplementary Fig. 21). In state 1, the predominant state in our substrate-trapped structure, the SpaR loop generates a narrow path, ~6Å in width, to make way for the substrate on its passage to the more spacious atrium. Stabilized by the formation of an antiparallel β-sheet, the hydrophobic side chain of SpaR Ile114 is exposed towards the channel lumen where it directly faces the translocating substrate (Fig. 5a and Supplementary Fig. 21). In state 2, no secondary structure elements can be observed and SpaR Ile114 is rotated away from the channel, which increases the width of the translocation path from ~6Å to approximately 10Å (Fig. 5a). PISA interface analysis revealed that neither of the two SpaR loop conformations forms stable interactions with residues building the translocation channel (state 1: ΔG: −4.4 kcal/mol, *P* = 0.86; state 2: ΔG: −4.1 kcal/mol, *P* = 0.85), demonstrating that the SpaR loop is a moveable element that enjoys conformational flexibility during substrate transport (Fig. 5a and Supplementary Table 2).

**Fig. 5.**
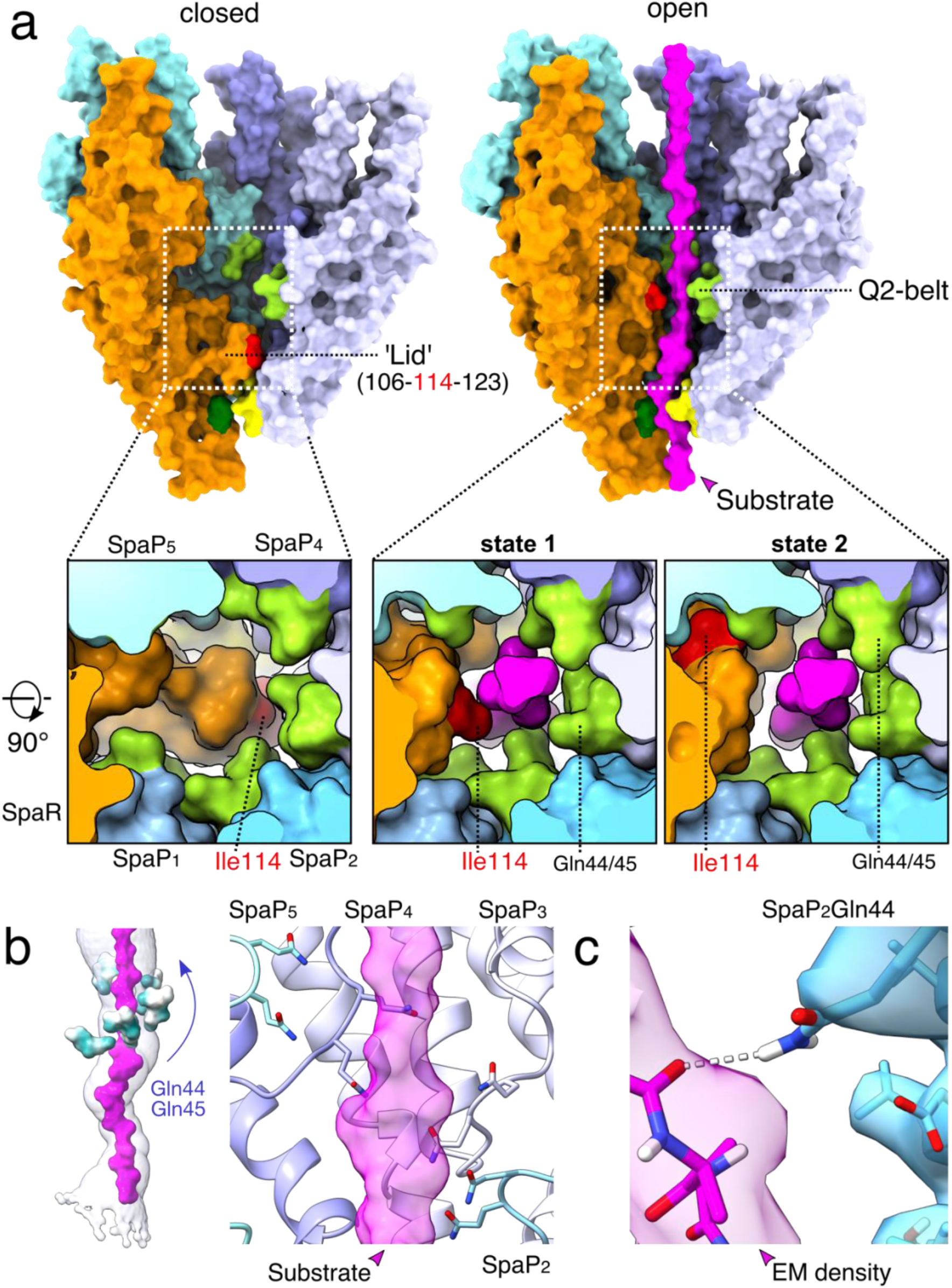
Conserved glutamines in SpaP and the SpaR lid together orchestrate substrate transport through the Q2-belt of the EA. **a**, Surface representations of the EA showing the SpaR loop/lid and surrounding Q2 belt. Q2 belt residues are shown in light green, M-gate residues in yellow. Numbers correspond to residues building the SpaR lid. The M-gate, Phe212 and Ile114 are displayed in yellow, dark green and red, respectively. **b**, Left: surface representations of the Q2 belt residues Gln44/45 colored by hydrophobicity. Right: close up of Q2 belt with Gln44/45 displayed in stick representation and SpaPs as ribbon diagrams. **c**, Hydrogen-bond formation between the substrate backbone and SpaP_2_ Gln45. The surfaces represent EM density (threshold: 0.016).

The area surrounding the substrate on the height of the upfolded SpaR loop is shaped by a loop which connects α-helices α2 and α3 in each of the five spirally-organized SpaP protomers (Fig. 5b). Reminiscent of the Q1-belt function, strictly conserved SpaP Gln44 and Gln45 localize within close proximity to the substrate, together generating an extended hydrogen-bonding donor/acceptor interface to engage with the backbone and polar side chains of the SptP substrate as it emerges from the M-gate and passes SpaR Ile114 (Fig. 5c and Supplementary Figs. 14 and 15). Because of the striking similarity with the Q1-belt, we decided to name this region of the translocation path the Q2-belt.

Together, it appears conceivable that the mobile SpaR loop functions akin to a linear ratchet in which Ile114 represents the ‘pawl’, whose conformational changes are physically triggered by the side chains or ‘teeth’ of the translocating substrate to support its unidirectional movement towards the atrium. Noteworthy, in state 2 (~10Å) but not state 1 (~6Å) of the SpaR loop, the Q2-belt provides sufficient space to also accommodate alpha helices, suggesting that the events that cause conformational switching of the SpaR loop may not be limited to unfolded substrates but may also be caused by translocating α-helices, which may provide an additional explanation for the evident ambiguity of the substrate density in our map.

Besides its role as a translocon, the EA also functions as a structural scaffold onto which the inner rod protein PrgJ assembles (Figs. 1c and 2a). Six PrgJ protomers interface with 1+5 SpaR+P proteins and previously unresolved lipids, together creating the atrium. This spacious chamber connects the channel, defined by the conical architecture of the EA, with the tunnel of the helical needle filament (Fig. 2). The novel lipids reside in a circular gap present in the upper EA, where they function to accommodate SpaP alpha helix α1 and stabilize the helical fork of PrgJ, together forming a nucleation seed that drives polymerization of the needle filament (Fig. 6a). All PrgJ proteins cross the EA/InvG gap to engage into β-strand complementation interactions formed between their N-termini and β-sheet β6 of the surrounding InvG subunits together stabilizing the inner rod within the confinement of the basal body (Supplementary Fig. 22). Notably, the lowest monomer PrgJ1 has a unique fold which extends horizontally, interfacing the SpaR loop connecting α-helices α2 and α3 before traversing SpaP1 to then cross the gap between the EA and the basal body clarifying earlier reports in which this part could not be resolved and PrgJ2 was speculated to interface in alternate locations (Supplementary Fig. 22)^6^. Likewise, also N-termini of previously unresolved PrgI1,4-5 protomers cross the EA/InvG gap to interact with SpaP, PrgJ, and the basal body component InvG, with each of these PrgI N-termini forming unique, plastic interactions with its respective environment, providing a rationale as to why some mutations localizing to the PrgI N-terminus abrogate filament assembly *in cellulo* but not *in vitro* (Supplementary Fig. 23)^24^.

**Fig. 6.**
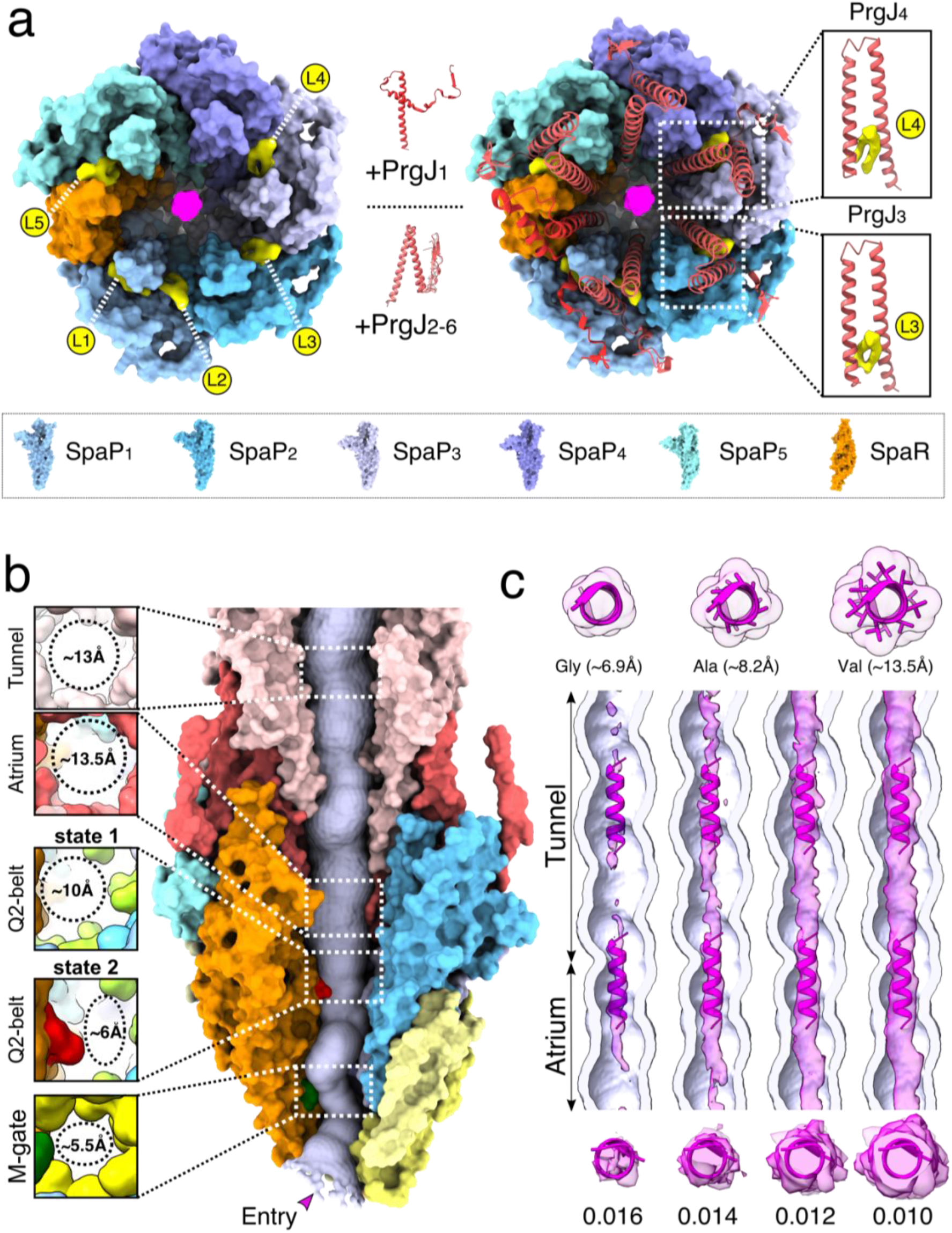
Density in the lipid-stabilized atrium and the filament tunnel suggests possible transport of partially folded substrates. **a**, Top views of EM densities corresponding to lipids (yellow, L1-L5) residing in a circular gap formed by SpaP and SpaR proteins in the EA. The lipids stabilize PrgJ α-helices α-1 and α-2 forming the PrgJ forks (right boxes; PrgJs displayed as red ribbon diagrams). **b**, Size constraints of the substrate translocation path through the needle complex. M-gate (ø ~5.5Å), Q2-belt (ø ~6Å, state 1; ~10Å, state 2), atrium (ø ~13.5Å), tunnel (ø ~13Å). **c**, Top: example α-helices and surface diameter calculations of poly-Gly, poly-Ala and poly-Val α-helices. Middle: vertical cross sections through the filament tunnel with density corresponding to the SptP3x-GFP substrate shown at increasing map thresholds. Bottom: secondary structure elements illustrate that the substrate EM density at high thresholds is sufficiently bulky to accommodate α-helices.

Further, the needle complex structure presented here provides a refined view on the helical PrgI filament which was built *de novo* into our non-symmetrized C1 map and hence, unlike in all other structures, no helical symmetry or restraints have been imposed. With an average axial helical rise of 4.41Å between subunits and a pitch of ~5.5 subunits, our C1 filament grows ~24.3Å per turn compared to ~23.8Å (6dwb), ~23.3Å (6ofh) and ~23.1Å (2lpz) in models obtained by helical reconstruction cryo-EM and NMR, respectively^7,8,24^. The quality of our map allowed us to model 72 PrgI subunits, covering a distance of ~36 nm and therefore our filament accumulates a total size difference of at least ~5.5Å. Despite this difference, the filament tunnel of our substrate-engaged structure adopts the form of a right-handed helix with a minimal inner diameter of ~13Å, which is indistinguishable from published apo-state structures and sufficiently large to accommodate α-helices (Fig. 6b)^7,8,14,24^.

At the passage from the Q2-belt to the atrium, the density corresponding to the substrate diminishes, demonstrating that, in contrast to the three tight interfaces seen in the EA, the wider lumen in the atrium (~13.5Å versus ~5.5-10Å) provides the substrate with higher conformational flexibility (Fig. 2 and 6b,c and Supplementary Fig. 6). Interestingly, the substrate density reappears in the upper atrium from where it continues through the tunnel of the filament (Fig. 6c). Here, it assumes a tubular shape which, especially at higher map thresholds, is remarkably similar to those of α-helices at low resolution, indicating that substrates that enter the secretion system potentially retain secondary structure elements or, alternatively, may partly refold during their passage through the filament (Fig. 6c).

## Discussion

Here, we report the first high-resolution snapshot of a type-III secretion system in an active, substrate-engaged conformation, providing insight into the molecular basis of protein transport across the bacterial envelope, a process that is fundamental to the virulence of many pathogenic bacteria and the motility of all flagellated bacteria. Our substrate-engaged structure reveals the complete secretion channel through the EA core complex, confirming its role as an entry portal to the needle complex.

Surprisingly, the EA lumen exhibits most of the conformational changes seen in the substrate-engaged structure, an unexpected finding given the sheer size and complexity of the needle complex machine. This agrees with, albeit at lower resolutions, our earlier structure and with visualizations of *in situ* needle complexes contacting host cells, together suggesting that the needle complex forms a largely static channel, in contrast to other more dynamic secretion machines^18,9,25^. In fact, many residues involved in substrate translocation localize to loop regions, which together with low rmsd values between apo-state and substrate-engaged structures, supports the concept that the basal body rings and the bulk of the SpaPQR complex provide a scaffold to position critical residues in the secretion channel. It appears plausible that this rigid architecture is a necessity to traverse the bacterial envelope, to provide a stable docking base for the dynamic components of the cytoplasmic sorting platform and simply to withstand the forces two moving cells but also the translocation process itself exert on the secretion system^4,26,27^.

In line with this concept, all six PrgJ proteins of the inner rod and PrgI2,4 of the filament base engage into interactions with the surrounding InvG proteins, together stabilizing the translocation conduit within the confinement of the basal body. To connect the conical shape of the EA with the helical filament, the inner rod protein PrgJ assumes a fold of a helical fork that is similar to those of the filament protein PrgI (Supplementary Figs. 22 and 23). Consequently, lipids that we find determine the shape of PrgJ, and hence the entire rod, are vital to its function as a nucleation seed that facilitates needle polymerization.

Strikingly, our substrate-trapped structure confirms the essential role of the EA core complex in T3 secretion, forming a conserved three-point interface with the substrate. The EA channel contains two hydrophilic Q-belts sandwiching the hydrophobic M-gate and movable SpaR lid, which together function as a gate and guide, to engage and steer substrates to the filament. By establishing complementary hydrophilic interactions with the substrate backbone, conserved glutamines facilitate effector protein translocation in a side chain-independent manner, an intriguing possibility due to the variety of different substrates accepted by T3SSs, which has also been exploited biotechnologically for secretion of artificially designed substrates (e.g. nanobodies, DARPins, monobodies, spider silk monomers, and viral epitopes)^28–30^.

Our data confirms the M-gate forms the main constriction in apo-state structures that shifts just enough to allow substrate passage. Evidently, this gate functions in flagellar systems as a seal to prevent unwanted leakage of metabolites and mutations increasing the gate size in our system impede cellular growth and T3SS secretion, which we believe is a result of creating leaky pores in the inner membrane, that likely impede the proton-motive-force (PMF) believed to power T3SS secretion^23,31^.

The unprecedented resolution achieved here enabled us to decipher a conformational switch of the SpaR lid which first folds up to make way for the translocating substrate to then function akin to a linear ratchet to facilitate continuous unidirectional motion of the unfolded substrate towards the filament. In closed states, the conformation of the SpaR loop can be traced only with sufficiently smoothed maps and at high map thresholds in both: isolated needle complexes (our structure & 6pep) and in the recombinant EA from *S. flexneri* (6r6b), suggesting that SpaR Leu110 and Ile114 are only weakly associated with the M-gate located below^6,12^. Interestingly, the flagellar homologue FliR typically utilizes two conserved phenylalanines, which tightly pack into the hydrophobic methionine network, a structural adaptation likely necessary to withstand the centrifugal force caused by the rotating flagellum and hence to maintain bacterial integrity (Supplementary Fig. 16)^15,12,13^.

Our substrate-engaged structure reveals continuous density throughout the needle complex conduit, corresponding to mostly non-folded SptP3x-GFP sequences in the EA channel. In fact, the N-termini of many substrates are heterogenous in sequence and predicted to be intrinsically disordered over a length of ~15-35 residues and therefore are sufficiently long to consecutively penetrate through all three constrictions, which we envision could be the most important unifying feature of T3SS signal sequences^20^. In favor of this concept, chaperones rather than N-terminal signal sequences were found to be crucial for routing effector proteins to the injectisome sorting platform^4^. Conceivably, it appears tempting to assume that effector proteins experience a selection pressure that drives the evolution of their N-termini towards non-folding sequences to facilitate their loading into the needle complex and hence ensure secretion.

However, SicP chaperone proteins maintain the chaperone-binding domain of SptP in a non-globular state containing α-helices, which raises the intriguing question whether or not further unfolding prior to entry into the secretion system is required^3^. Evidently, for α-helices to be directly translocated through the EA channel, the narrow, ~5.5Å-wide lumen of the M-gate would have to open further (Fig. 6b,c). While we cannot rule out that α-helices are in fact directly translocated through the EA, the tubular appearance of the density corresponding to the substrate in the filament in our map suggests that α-helices may fold during their passage through the needle complex. Either way, the transport of α-helices is likely advantageous, accelerating the folding of effector proteins upon arrival in the host cell cytoplasm and allowing them to elicit their effector functions faster, a process that would benefit pathogen fitness. Also, helical filament assembly could be more effectively achieved by docking of the readily formed helices into their cognate interfaces at the distal end of the growing filament.

Future studies will also have to address the question whether the constrictions inside the EA channel are passive components that only change their conformation as a consequence of the penetrating substrate or, alternatively, are regulated by active gating mechanisms. Given that hardly any conformational differences are seen in the scaffolding parts of the SpaPQR proteins (Supplementary Fig. 9), it is difficult to imagine a cytoplasmic signal translating up to the M-gate or SpaR ‘lid’ prior to substrate engagement, instead suggesting that these structural elements shift in response to the substrate physically travelling through the channel. However, we cannot rule out the possibility that gating of the EA portal occurs through SpaS, which is mostly absent in our complexes but likely orchestrates the shape of the SpaQ loops building the Q1-belt in native injectisomes. Alternatively, SpaS could represent a physical link providing guidance for the approaching disordered N-terminus to help finding its way into the narrow cytoplasmic opening of the EA portal.

Based on the structure and biochemical work presented here, we propose a model of substrate secretion through the T3SS export apparatus (Fig. 7). 1) After chaperone removal and substrate unfolding by the ATPase, the substrate is guided to the EA portal engaging with the Q1-belt which accommodates effector proteins independent from their sequence. 2) The substrate transports up through the methionine network, disrupting its hydrophobic interface causing the methionines to shift apart and opens the M-gate. 3) The SpaR loop, resting above the M-gate and blocking the channel, extends upwards, assuming an open conformation, which together with the surrounding Q2-belt creates an interface to engage the substrate. 4) The upfolded SpaR loop contacts the passing substrate, functioning similar to a linear ratchet to prevent a back slipping of the substrate and together with the Q2 belt guides the substrate up through the atrium and into the lumen of the filament.

**Fig. 7.**
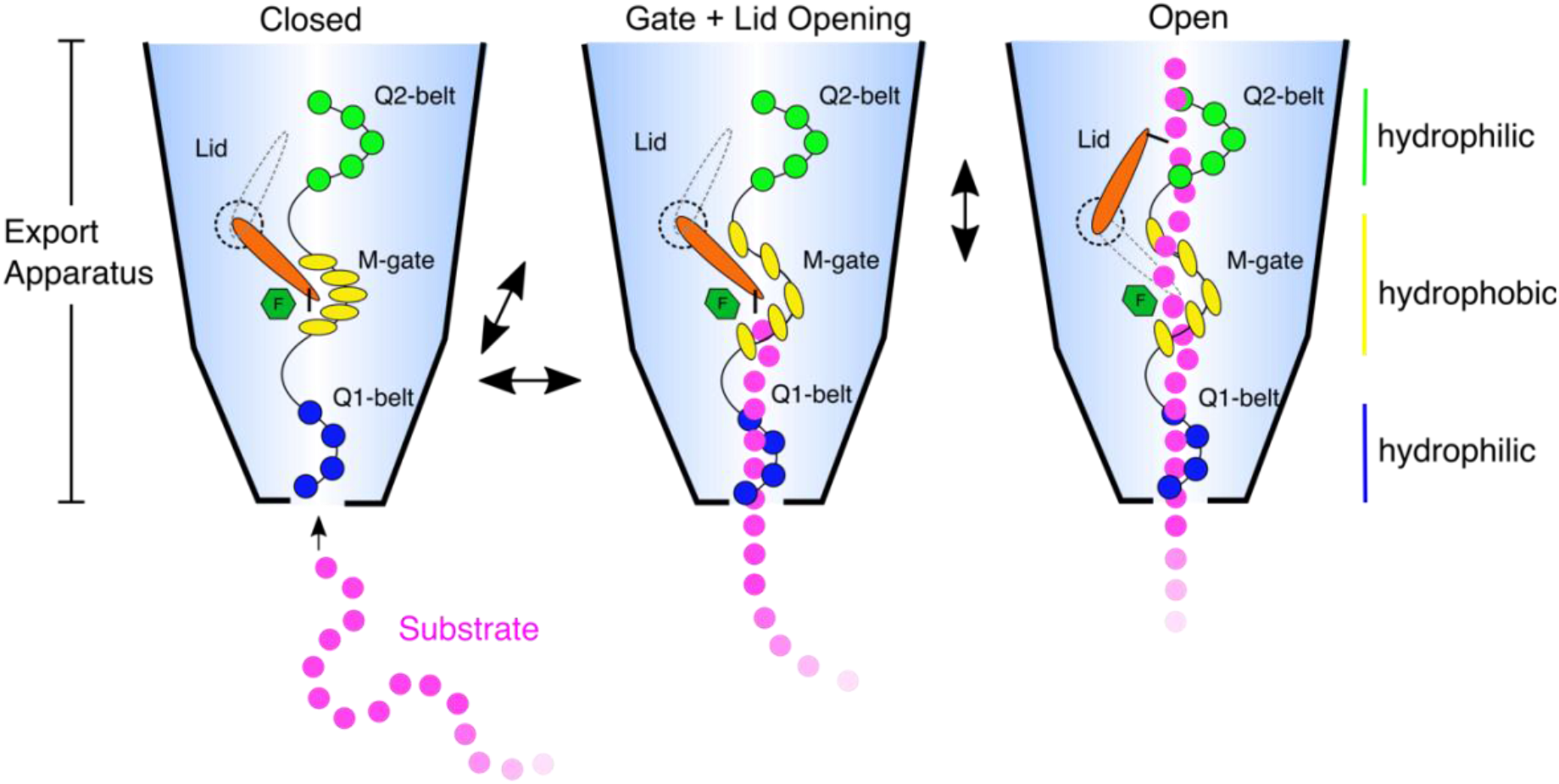
Model for substrate translocation through the EA of T3SSs. Disordered N-termini of effector proteins (magenta circles) enter the Q1-belt (blue circles) of the EA facing the bacterial cytoplasm. Successive binding of the substrate backbone to conserved glutamines facilitates penetration through the Q1-belt and loading of the substrate into the needle complex, a process putatively fueled by the concerted action of the ATPase (InvC) and the proton motive force (PMF). The substrate then penetrates the M-gate. Disruption of its hydrophobic interface (Met186-Met187: yellow ovals; SpaR Phe212: green hexagon) expands the gate and causes the SpaR loop (orange ratchet) to flip into a vertical position together generating a narrow path for the translocation of the unfolded substrate chain. The substrate then proceeds up to the Q2-belt (green circles) and SpaR Ile114 (black pawl) which together stabilize the substrate above the M-gate and the loop likely acts as a linear ratchet to engage and steer substrates further towards the atrium and filament, ultimately facilitating effector protein secretion.

## Methods

### Bacterial strains and plasmids

Experiments were conducted using a *S. enterica* sv. Typhimurium non-flagellated strain SB905 carrying the T3SS transcriptional regulator gene *hilA* expressed on the pSB3291 plasmid using an araBAD promoter as previously described^18^. The SptP-based substrate and the SptP chaperone sicP, were both expressed using a pACYCDuet-1 CmR plasmid (Merck Chemicals). The substrate construct consists of an N-terminal signal sequence, a SicP-chaperone-binding domain, three SptP effector domain repeats fused to eGFP and a 3× FLAG tag^18^.

### Bacterial Growth Assay

Overnight cultures of *Salmonella* SB905 SpaP knockout strains complemented with either WT SpaP or with GGG, AAA, CCC, WWW, and FFF substitutions were grown with or without 0.3 M NaCl supplementation (injectisome inducing vs non inducing conditions) under antibiotic selection. The following morning, cultures were normalized to an OD_600_ of 0.5 before diluting 1:10 in either non-inducing LB, or inducing media, LB supplemented with 0.3 M NaCl and 0.012% w/v arabinose, for injectisome assembly. Cultures were sampled at 1 hour intervals and OD_600_ measurements were recorded using a spectrophotometer to calculate growth curves. Three independent experiments of three biological replicates were used for each strain and condition. Data and statistical analyses were visualized using the GraphPad Prism8 software package. Needle Complex Counting

Overnight cultures of *Salmonella* SB905 SpaP knockout strains complemented with either WT SpaP or with M-gate substitutions were grown with 0.3 M NaCl supplementation under antibiotic selection. The following morning, cultures were diluted 1:10 in injectisome-inducing media, LB supplemented with 0.3 M NaCl, 0.012% w/v arabinose and without antibiotics. After 5 hours of growth, cultures were diluted to an OD_600_ of 1.0 and 1 ml was collected and centrifuged for 5 min at RT, approx. 16000 x *g.* The pellets were resuspended in approximately 30 μl of LB and incubated at RT for 5 min. 1 ml of ice-cold water was mixed with the cells which were then incubated on ice for 5 min. 1.5 μl of DNAse (Thermo Fisher Scientific) was added along with 20 μl of 1 M MgCl2 and incubated at RT for 10 min. 40 μl of 0.5 M EDTA and 150 μl of 1 M Tris-Cl (pH 7.5) were added and the cells were centrifuged for 30 min at 4°C, approx. 16000 x *g*. Pellets were resuspended with 1 ml of ice cold TE buffer and centrifuged again. Pellets were then resuspended in 10-50 μl of ice cold TE buffer and incubated at 4°C for approximately 20 min while shaking. Cells were applied to TEM grids and stained (see below) before imaging with a Thermo Fisher Talos L120C TEM. Twenty cells were imaged at varying magnifications and needle complexes were counted by three colleagues, independently. Counts from each person for each cell were averaged and the data was visualized using the GraphPad Prism8 software package. Purification of needle complexes

Needle complexes were purified either from *Salmonella* SB905 WT or SpaP KO strains as described previously^5,18^. Briefly, day cultures were grown for 5 hours in LB media supplemented with 0.3 M NaCl, 0.012% w/v arabinose and without antibiotics prior to harvesting. Needle complexes were extracted from the membranes using 0.4% w/v LDAO, and separated by CsCl centrifugation. Fractions containing assembled needle complexes were concentrated and used for negative staining or cryoEM. For purification of substrate-trapped complexes, SB905 expressing hilA and the substrate SptP3x-GFP with chaperone sicP was grown for 4 hours with an additional 2 hour induction period using 1 mM IPTG prior to harvesting. To enrich for substrate-trapped complexes, CsCl fractions containing needle complexes and the substrate were pooled and incubated with Anti-FLAG M2 magnetic beads (M8823, Sigma Aldrich/Merck KGaA) prewashed in FR3 buffer (10 mM Tris-HCl, pH 8.0, 0.5 M NaCl, 5 mM EDTA, 0.1% w/v LDAO) for 2.5 hours under gentle agitation at 4°C. The beads were subsequently washed 12 times with FR3 buffer for 10 min each. Beads were eluted with FR3 buffer twice as mock elutions and subsequently eluted twice for 45 min with 2 mg/ml 3xFLAG peptide in FR3 buffer. These elutions were then pooled and pelleted in a Beckman TLA-110 rotor at 90k rpm for 35 min. T3SS pellets were resuspended for at least 1 hour in FR3 buffer under agitation at 4°C before TEM imaging. Negative staining TEM 4 μl of diluted sample was applied to carbon-coated copper grids and incubated for 40 seconds. Grids were glow discharged before for 30 seconds at 25 mA using a GloQube^®^ Plus Glow Discharge System (Electron Microscopy Sciences). The sample was blotted off, and the grid was washed briefly with 4 μl of staining solution (2% w/v PTA, adjusted to pH 7.0 with NaOH) and then stained with 4 μl of the staining solution for 20 sec. The stain was blotted off and the grids were air-dried for at least 1 min. Grids were imaged using a Thermo Fisher Scientific Talos L120C TEM with a 4K Ceta CEMOS camera.

### SPA cryoEM sample preparation and data collection

Purified apo-state needle complexes were applied to Quantifoil grids with either an additional layer of amorphous carbon (<1.6nm thick) or graphene oxide. Purified, substrate-trapped needle complexes were applied to Quantifoil grids floated with an approx. 1.1 nm layer of amorphous carbon on top. 4 μl of sample was applied onto glow-discharged grids (30 sec, 25 mA) and allowed to disperse for 0.5-2 min. The grids were blotted for 4-7 sec set at 100% humidity and plunge frozen in a liquid propane/ethane mixture cooled to liquid nitrogen temperatures, using a Thermo Fisher Scientific Vitrobot Mark V. Vitrified samples were imaged on a Thermo Fisher Scientific Titan Krios TEM operating at 300 kV and equipped with a field emission gun (XFEG) and a Gatan Bioquantum energy filter. Movies consisting of 25 (apo) or 50 (substrate-trapped) frames, were automatically recorded using Thermo Fisher Scientific EPU software and a Gatan K2 or K3 camera, at 0.3 – 5.2 μm defocus in counting mode (Supplementary Table 1).

### SPA image processing

SPA was performed using Relion 3.0 for the apo state needle complex and Relion 3.1-beta for the substrate-trapped needle complex^32,33^. Movies were motion-corrected^34^, dose-weighted and the CTF was determined using CTFFIND4^35^. Particles were automatically picked from the motion– corrected micrographs using crYOLO^36^ trained with a subset of manually picked particles. Particles were extracted and binned for several rounds of 2D classification. A cleaned and unbinned data set was obtained by re-extraction and aligned to a rotationally averaged structure. Focused refinements with and without applying symmetry were performed to the individual substructures using respective 3D masks. After converged refinements, per particle CTF and Bayesian polishing was used to generate a new data set for another round of focused refinements. Final rounds of refinement were performed without any masking of particles. Overall gold-standard resolution (Fourier shell correlation (FSC) = 0.143) and local resolution as well as sharpened maps (B-factor: −30) were calculated with Relion 3.1-beta^32^.

### Model building, refinement and validation

#### Apo state

Model building into the T3SS apo-state map started by placing homology models for SpaP, SpaQ and SpaR, which were generated with SWISS-MODEL using *Vibrio mimicus* FliP, FliQ and FliR structures as templates (6s3l); PrgJ was modelled using Phyre2^13,37,38^. Together with PrgI (2lpz), homology models for SpaP, SpaQ, SpaR and PrgJ were first fitted into the EM map using the fit-in-map tool in USCF Chimera (v1.14) and then manually extended with Coot (v0.9-beta)^7,39,40^. A first refinement was performed with Rosetta controlled via StarMap (manuscript in preparation), followed by interactive refinement against the map density with ISOLDE (v.1.0b5), a molecular dynamics-guided structure refinement tool within ChimeraX (v.0.93)^41,42^. The resulting coordinate file was further refined with Phenix.real_space_refine (v.1.18-6831) using reference model restraints, strict rotamer matching and disabled grid search. Model validation was carried out using MolProbity server and EMRinger within the Phenix software package (Supplementary Table 1)^43–46^.

#### Substrate-trapped state

Existing models of PrgH (3gr1), PrgK (3gr5), InvG (4g08, G34-I173: lower OR), SpaP, SpaR, SpaS, PrgJ and PrgI were rigid body-fitted into the electron density map using the fit-in-map tool in UCSF Chimera (v1.14), followed by manual rebuilding in Coot (v0.9-beta)^39,40,47,48^. The upper OR (InvG: E174-G557) was first built into a C15-symmetrized and focus-refined map using Coot (v0.9-beta) for *ab-initio* model building, followed by rigid body-fitting into the C1 density map using the fit-in-map tool in UCSF Chimera v1.14^39,40^. Interactive refinement against the C1 map density was performed with ISOLDE (v.1.0b5)^41^. The resulting coordinate file was further refined with Phenix.real_space_refine (v.1.18-6831) using reference model restraints, strict rotamer matching and disabled grid search^45^. Model validation was carried out using MolProbity server and EMRinger (Supplementary Table 1)^43,44,46^. The translocation channel through the export apparatus and lumen of the needle filament was calculated using HOLE^49^. The helical rise of the PrgI filaments from the substrate-trapped filament, 2lpz, 6ofh, and 6dwb were measured by running the Rosetta tool make_symmdef_file.pl for each consecutive pair of PrgI protomers in the direction from base to tip^7,8,24,50^. Measurements were limited to 26 protomers and started from PrgI12 in the substrate-trapped filament. UCSF Chimera and ChimeraX were used for molecular visualization^39,42^.

## Acknowledgements

We thank all members of the Marlovits Laboratory including Catalin Bunduc, Biao Yuan, and Barbara Grueter, for their support of this project. We would like to thank Wolfgang Lugmayr and Frank DiMaio for their help with StarMap, Rosetta, and the filament analysis. We would also like to thank Tristan Croll for his significant support with ISOLDE. High-performance computing was possible through access to the HPC at DESY/Hamburg (Germany) and the Vienna Scientific Cluster (Austria). Part of this work was performed at the CryoEM Facility at CSSB, supported by the UHH and DFG grant numbers (INST152/772-1 | 152/774-1 | 152/775-1 | 152/776-1 | 152/777-1 FUGG).

## Author contributions

SM, DF designed experiments. SM, DF generated constructs. NGM and MP generated knockout strains. SM, VK, OV purified complexes. SM, DF performed biochemical assays. SM, JW vitrified samples and collected cryoEM images. SM collected negative stain images. SM, DF, NGM built the atomic model. SM DF, NGM, TCM interpreted data. TCM processed cryoEM data. SM, DF, TCM wrote and revised the paper. All authors read, corrected and approved the manuscript. TCM conceived the study and supervised the project.

## Data and materials availability

maps have been deposited at the EMDB database. Models have been deposited at the PDB.

## References

1. Hueck, C. J. Type III protein secretion systems in bacterial pathogens of animals and plants. Microbiol. Mol. Biol. Rev. MMBR 62, 379–433 (1998).

2. Galán, J. E. & Waksman, G. Protein-injection machines in bacteria. Cell 172, 1306–1318 (2018).

3. Stebbins, C. E. & Galán, J. E. Maintenance of an unfolded polypeptide by a cognate chaperone in bacterial type III secretion. Nature 414, 77–81 (2001).

4. Lara-Tejero, M., Kato, J., Wagner, S., Liu, X. & Galán, J. E. A sorting platform determines the order of protein secretion in bacterial type III systems. Science 331, 1188–1191 (2011).

5. Schraidt et al. Topology and organization of the *Salmonella* typhimurium type III secretion needle complex components. PLoS Pathog. 6, e1000824 (2010).

6. Hu, J. et al. T3S injectisome needle complex structures in four distinct states reveal the basis of membrane coupling and assembly. Nat. Microbiol. 2010-2019 (2019) doi:10.1038/s41564-019-0545-z.

7. Loquet, A., Sgourakis, N. G., Gupta, R. & Giller, K. Atomic model of the type III secretion system needle. Nature 486, 276–279 (2012).

8. Hu, J. et al. Cryo-EM analysis of the T3S injectisome reveals the structure of the needle and open secretin. Nat. Commun. 9, 3840–3840 (2018).

9. Park, D. et al. Visualization of the type III secretion mediated *Salmonella-host* cell interface using cryo-electron tomography. eLife 7, e39514 (2018).

10. Kubori, T. et al. Supramolecular structure of the *Salmonella* typhimurium type III protein secretion system. Science 280, 602–605 (1998).

11. Marlovits, T. C. et al. Structural insights into the assembly of the type III secretion needle complex. Science 306, 1040–1042 (2004).

12. Johnson, S., Kuhlen, L., Deme, J. C., Abrusci, P. & Lea, S. M. The structure of an injectisome export gate demonstrates conservation of architecture in the core export gate between flagellar and virulence type III secretion systems. mBio 10, e00818–19 (2019).

13. Kuhlen, L. et al. The substrate specificity switch FlhB assembles onto the export gate to regulate type three secretion. Nat. Commun. 11, 1–10 (2020).

14. Lunelli, M. et al. Cryo-EM structure of the *Shigella* type III needle complex. PLoS Pathog. 16, e1008263 (2020).

15. Kuhlen, L. et al. Structure of the core of the type III secretion system export apparatus. Nat. Struct. Mol. Biol. 25, 583–590 (2018).

16. Galán, J. E. & Wolf-Watz, H. Protein delivery into eukaryotic cells by type III secretion machines. Nature 444, 567–573 (2006).

17. Schlumberger, M. C. et al. Real-time imaging of type III secretion: *Salmonella* SipA injection into host cells. PNAS 102, 12548–12553 (2005).

18. Radics, J., Königsmaier, L. & Marlovits, T. C. Structure of a pathogenic type 3 secretion system in action. Nat. Struct. Mol. Biol. 21, 82–87 (2014).

19. Dohlich, K., Zumsteg, A. B., Goosmann, C. & Kolbe, M. A substrate-fusion protein is trapped inside the type III secretion system channel in *Shigella flexneri*. PLoSPathog 10, e1003881 (2014).

20. McDermott, J. E. et al. Minireview: Computational prediction of type III and IV secreted effectors in gram-negative bacteria. Infect. Immun. 79, 23–32 (2011).

21. Buchko, G. W. et al. A multi-pronged search for a common structural motif in the secretion signal of *Salmonella enterica* serovar Typhimurium type III effector proteins. Mol. Biosyst. 6, 2448–2458 (2010).

22. Kimbrough, T. G. & Miller, S. I. Contribution of *Salmonella* typhimurium type III secretion components to needle complex formation. PNAS 97, 11008–11013 (2000).

23. Ward, E. et al. Type-III secretion pore formed by flagellar protein FliP. Mol. Microbiol. 107, 94–103 (2018).

24. Guo, E. Z. et al. A polymorphic helix of a *Salmonella* needle protein relays signals defining distinct steps in type III secretion. PLOS Biol. 17, e3000351 (2019).

25. Basler, M., Pilhofer, M., Henderson, G. P., Jensen, G. J. & Mekalanos, J. J. Type VI secretion requires a dynamic contractile phage tail-like structure. Nature 483, 182–186 (2012).

26. Diepold, A. et al. A dynamic and adaptive network of cytosolic interactions governs protein export by the T3SS injectisome. Nat. Commun. 8, 15940 (2017).

27. Hu, B., Lara-Tejero, M., Kong, Q., Galán, J. E. & Liu, J. In situ molecular architecture of the *Salmonella* Type III secretion machine. Cell 168, 1065–1074 (2017).

28. Chabloz, A. et al. Salmonella-based platform for efficient delivery of functional binding proteins to the cytosol. Commun. Biol. 3, 1–11 (2020).

29. Widmaier, D. M. et al. Engineering the *Salmonella* type III secretion system to export spider silk monomers. Mol. Syst. Biol. 5, 309 (2009).

30. Rüssmann, H. et al. Delivery of epitopes by the *Salmonella* type III secretion system for vaccine development. Science 281, 565–568 (1998).

31. Paul, K., Erhardt, M., Hirano, T., Blair, D. F. & Hughes, K. T. Energy source of flagellar type III secretion. Nature 451, 489–492 (2008).

32. Zivanov, J., Nakane, T. & Scheres, S. H. W. Estimation of high-order aberrations and anisotropic magnification from cryo-EM data sets in RELION-3.1. IUCrJ 7, 253–267 (2020).

33. Zivanov, J. et al. New tools for automated high-resolution cryo-EM structure determination in RELION-3. eLife 7, e42166 (2018).

34. Zheng, S. Q. et al. MotionCor2: Anisotropic correction of beam-induced motion for improved cryoelectron microscopy. Nat. Methods 14, 331–332 (2017).

35. Rohou, A. & Grigorieff, N. CTFFIND4: Fast and accurate defocus estimation from electron micrographs. J. Struct. Biol. 192, 216–221 (2015).

36. Wagner, T. et al. SPHIRE-crYOLO is a fast and accurate fully automated particle picker for cryo-EM. Commun. Biol. 2, 1–13 (2019).

37. Bordoli, L. et al. Protein structure homology modeling using SWISS-MODEL workspace. Nat. Protoc. 4, 1–13 (2009).

38. Kelley, L. A., Mezulis, S., Yates, C. M., Wass, M. N. & Sternberg, M. J. E. The Phyre2 web portal for protein modeling, prediction and analysis. Nat. Protoc. 10, 845–858 (2015).

39. Pettersen, E. F. et al. UCSF Chimera - a visualization system for exploratory research and analysis. J. Comput. Chem. 25, 1605–1612 (2004).

40. Emsley, P., Lohkamp, B., Scott, W. G. & Cowtan, K. Features and development of Coot. Acta Crystallogr. D Biol. Crystallogr. 66, 486–501 (2010).

41. Croll, T. I. ISOLDE: A physically realistic environment for model building into low-resolution electron-density maps. Acta Crystallogr. Sect. Struct. Biol. 74, 519–530 (2018).

42. Goddard, T. D. et al. UCSF ChimeraX: meeting modern challenges in visualization and analysis. Protein Sci. 27, 14–25 (2018).

43. Barad, B. A. et al. EMRinger: Side-chain-directed model and map validation for 3D Electron Cryomicroscopy. Nat. Methods 12, 943–946 (2015).

44. Williams, C. J. et al. MolProbity: More and better reference data for improved all-atom structure validation. Protein Sci. Publ. Protein Soc. 27, 293–315 (2018).

45. Liebschner, D. et al. Macromolecular structure determination using X-rays, neutrons and electrons: Recent developments in Phenix. Acta Crystallogr. Sect. Struct. Biol. 75, 861–877 (2019).

46. Lang, P. T. et al. Automated electron-density sampling reveals widespread conformational polymorphism in proteins. Protein Sci. Publ. Protein Soc. 19, 1420–1431 (2010).

47. Spreter, T. et al. A conserved structural motif mediates formation of the periplasmic rings in the type III secretion system. Nat. Struct. Mol. Biol. 16, 468–476 (2009).

48. Bergeron, J. R. C. et al. A Refined Model of the Prototypical *Salmonella* SPI-1 T3SS Basal Body Reveals the Molecular Basis for Its Assembly. PLOS Pathog. 9, e1003307 (2013).

49. Smart, O. S., Neduvelil, J. G., Wang, X., Wallace, B. A. & Sansom, M. S. P. HOLE: A program for the analysis of the pore dimensions of ion channel structural models. J. Mol. Graph. 14, 354–360 (1996).

50. DiMaio, F., Leaver-Fay, A., Bradley, P., Baker, D. & André, I. Modeling symmetric macromolecular structures in Rosetta3. PLoS ONE 6, e20450 (2011).

51. Kucukelbir, A., Sigworth, F. J. & Tagare, H. D. Quantifying the local resolution of cryo-EM density maps. Nat. Methods 11, 63–65 (2014).

52. Pintilie, G. & Chiu, W. Comparison of Segger and other methods for segmentation and rigid-body docking of molecular components in cryo-EM density maps. Biopolymers 97, 742–760 (2012).

53. Pintilie, G., Zhang, J., Chiu, W. & Gossard, D. Identifying components in 3D density maps of protein nanomachines by multi-scale segmentation. IEEENIH Life Sci. Syst. Appl. Workshop IEEENIH Life Sci. Syst. Appl. Workshop 2009, 44–47 (2009).

54. Madeira, F. et al. The EMBL-EBI search and sequence analysis tools APIs in 2019. Nucleic Acids Res. 47, W636–W641 (2019).

55. Robert, X. & Gouet, P. Deciphering key features in protein structures with the new ENDscript server. Nucleic Acids Res. 42, W320–W324 (2014).

56. Krissinel, E. & Henrick, K. Inference of macromolecular assemblies from crystalline state. J. Mol. Biol. 372, 774–797 (2007).

