## Supplementary Information for "Substrate-engaged type III secretion system structures reveal gating mechanism for unfolded protein translocation"

### This document contains:

Supplementary Figures 1 – 23

Supplementary Tables 1 and 2

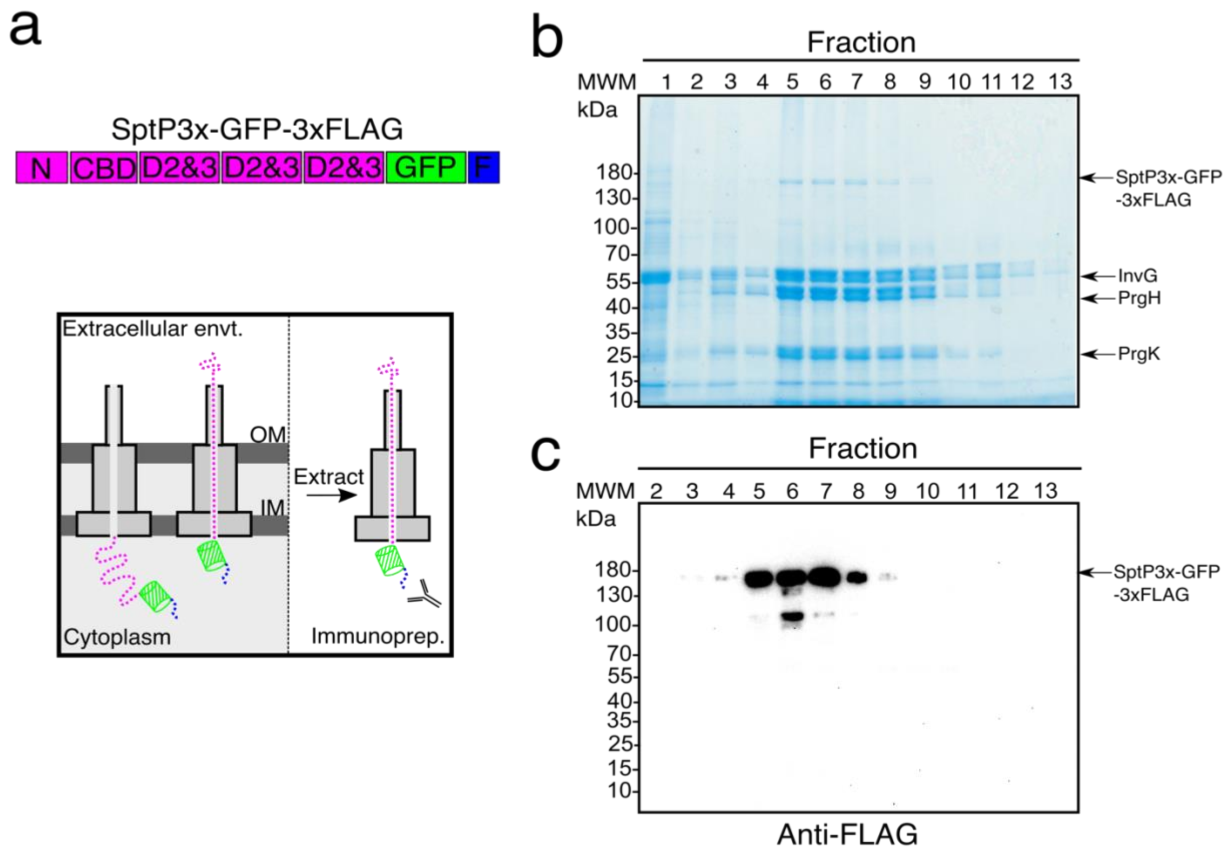


**Supplementary Figure 1: Needle complex purification. a**, Domain organization of the engineered SptP construct used in this study. N: N-terminal signal sequence; CBD: SicP-specific chaperone binding domain; D2&3: repeats of SptP effector domains 2 and 3; GFP: Unfolding resistant enhanced green fluorescent protein; F: 3xFLAG protein purification tag for pulldown experiments. Below panel: schematic on trapping substrates in needle complexes and subsequent immunoprecipitation. OM: outer membrane, IM: inner membrane. Substrate shown in magenta with a green cartoon representing GFP, and blue 3XFLAG tag. **b**, Coomassie-blue stain of CsCl gradient fractions obtained during needle complex purification. SptP3x-GFP migrates to its approximate molecular weight of 176 kDa. **c**, Western blot analysis of CsCl gradient fractions in (**b**) probed with an anti-FLAG antibody.

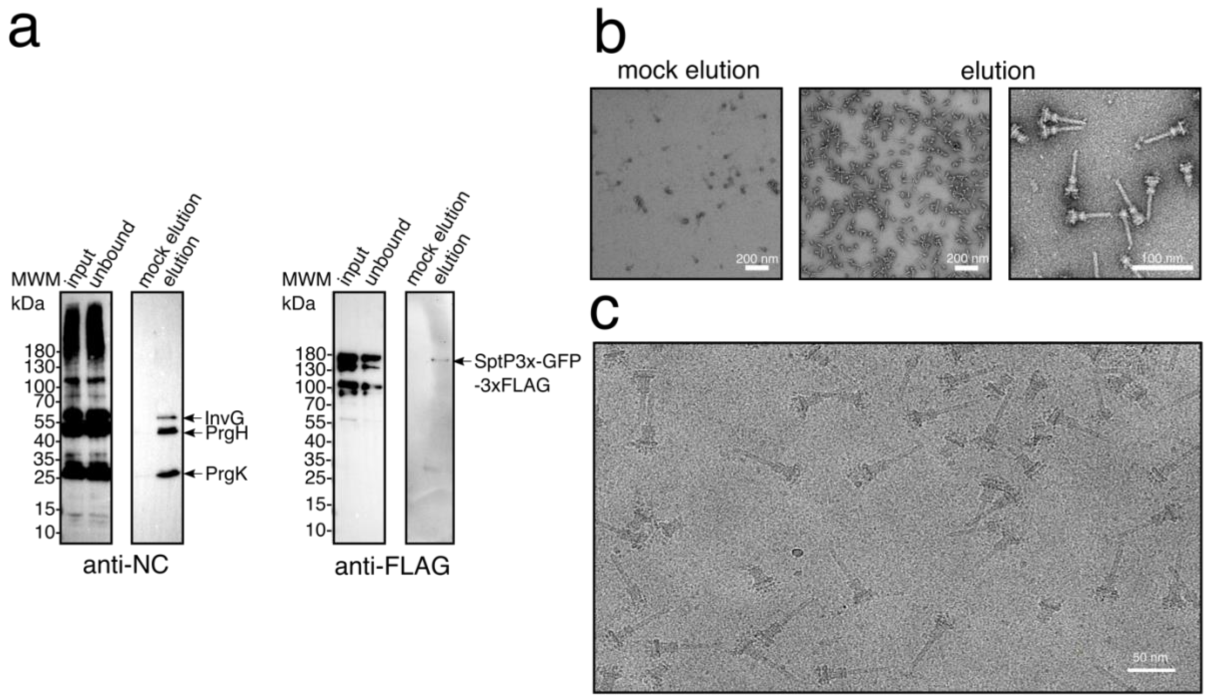


**Supplementary Figure 2: Immunoprecipitation of substrate-engaged needle complexes. a**, Western blot analysis of needle complex pulldown experiments using anti-FLAG magnetic beads probed with anti-needle complex antibodies recognizing inner (PrgH/K) and outer ring (InvG) proteins (left) and anti-FLAG antibody (right). Unbound: Supernatant of pulldown sample. Mock elution: Elution step with buffer lacking the FLAG peptide eluent. **b**, Negative-stain TEM of samples taken from mock (left) and FLAG peptide (right) elutions at two magnification levels. Scale bars: 100 nm and 200 nm. **c**, Representative micrograph of plunge frozen substrate-engaged needle complexes. Scale bar: 50 nm.

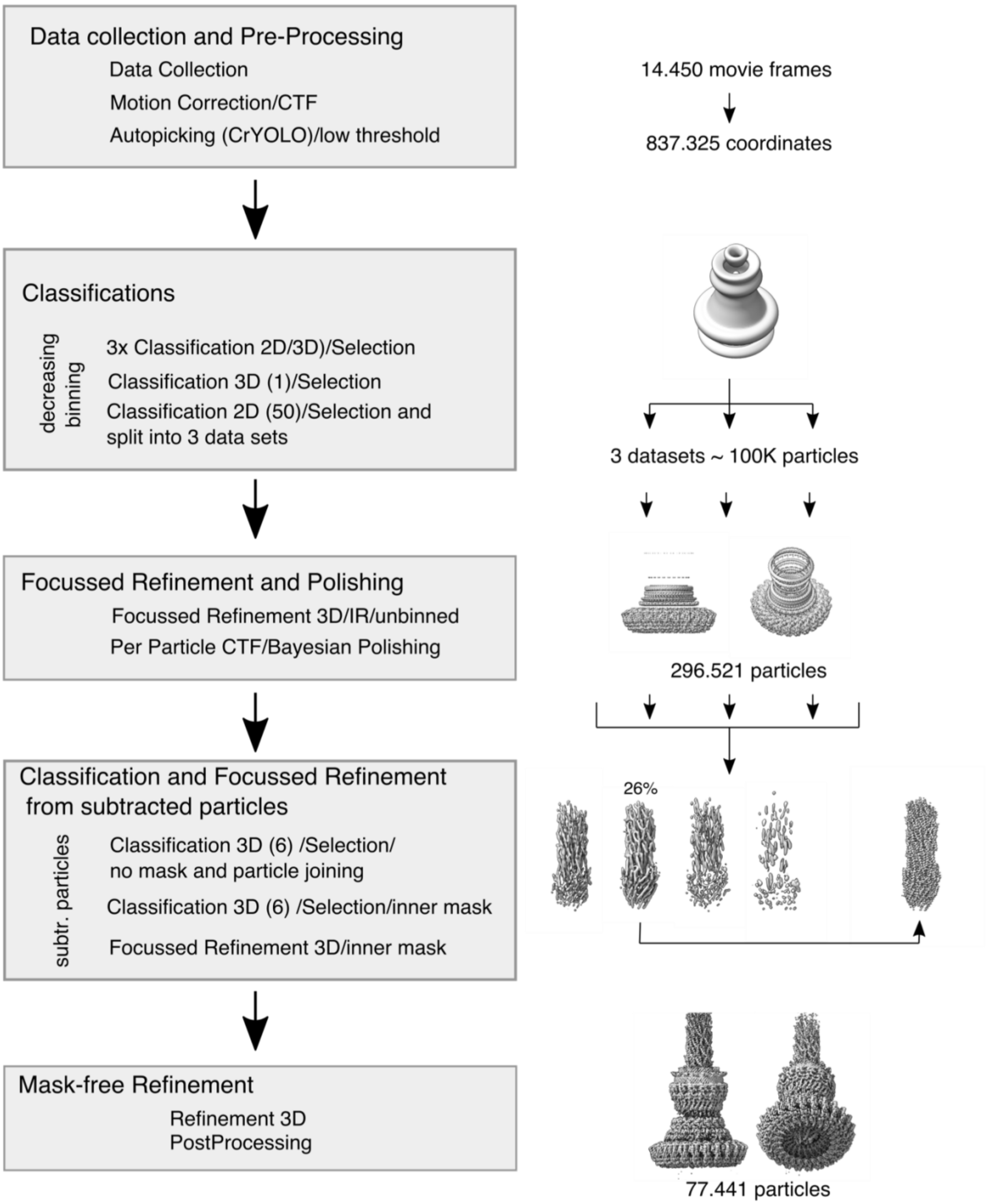


**Supplementary Figure 3: Cryo-EM data collection and single particle reconstruction procedure.**

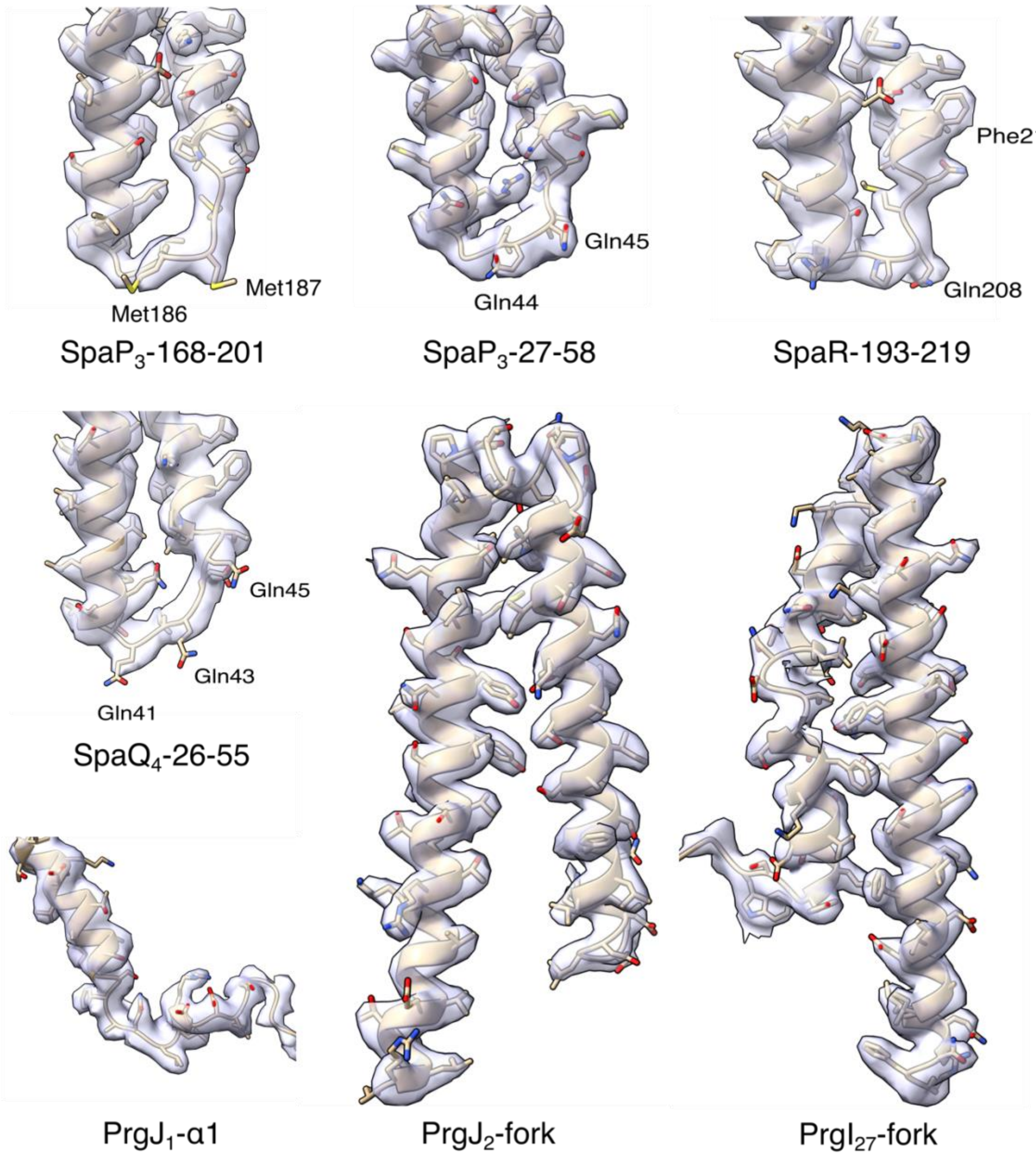


**Supplementary Figure 4: Example cryo-EM densities.** Shown are regions containing residues directly involved in substrate translocation through the EA and protein components building the inner rod (atrium) and filament (tunnel). Met186 and Met187 of SpaP$_3$ (M-gate), Gln44 and Gln45 of SpaP$_3$ (Q2-belt), Gln208 and Phe212 of SpaR (Q1-belt/M-gate) and Gln41, Gln43 and Gln45 of SpaQ$_4$ (Q1-belt). PrgJ$_1$ alpha helix 1 and the forks of PrgJ$_2$ and PrgI$_{27}$. Models are shown in ribbon cartoons with stick representations.

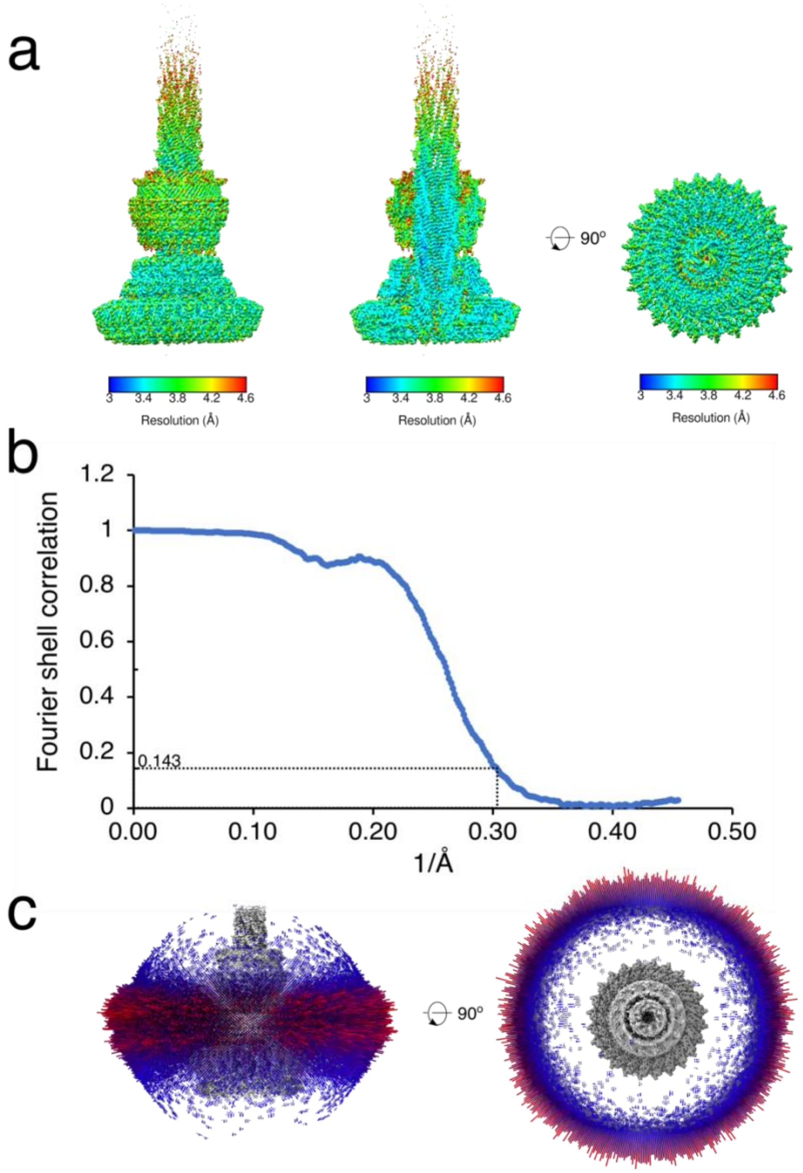


**Supplementary Figure 5: Single particle reconstruction of the substrate-engaged needle complex from *S. enterica* sv. Typhimurium. a**, Local resolution estimation of the masked C1 map (left), a vertical cross section (middle) and a top view (right) calculated by ResMap[51]. **b**, Fourier Shell Correlation (FSC) of the masked C1 map. Global resolution at the gold-standard FSC threshold of 0.143 is ~3.36Å. **c,** Side (left) and top (right) view of the angular distribution plot of all needle complex particles that contributed to the final C1 map. Height and color (from blue to red) of the cylinder bars is proportional to the number of particles.

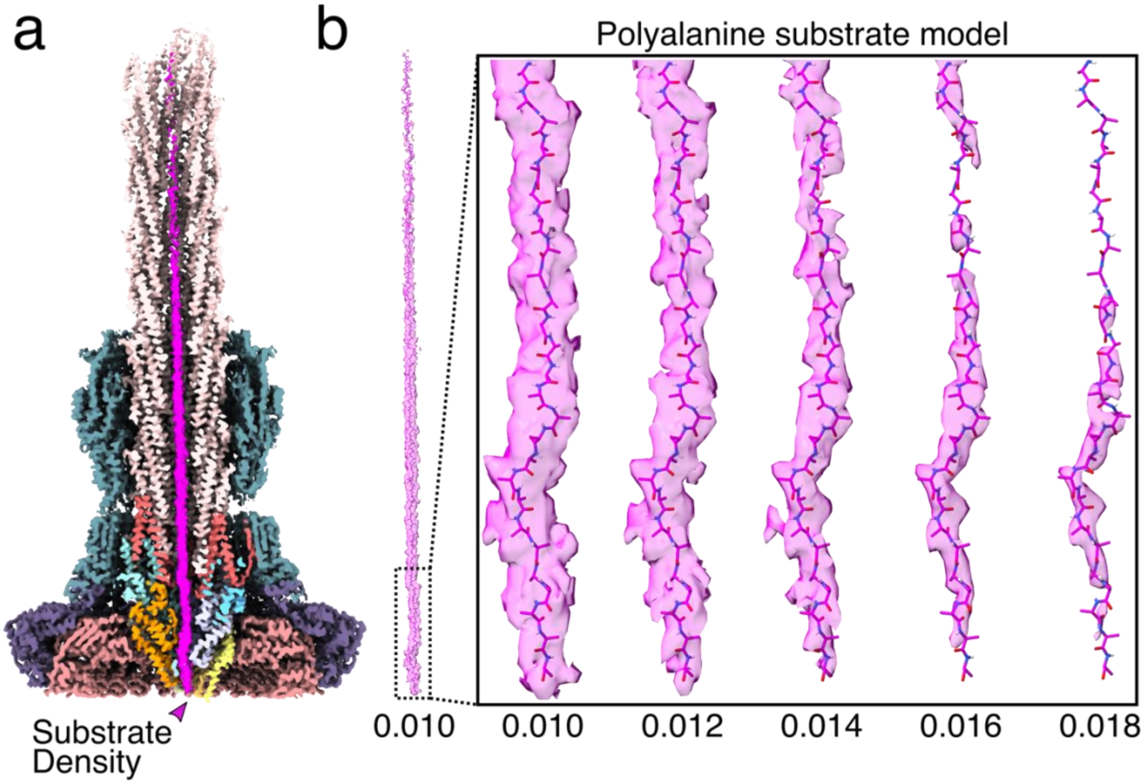


**Supplementary Figure 6: The SptP3x-GFP substrate adopts a non-globular conformation inside the needle complex translocation channel. a**, Cross section of the C1 map of the needle complex with the substrate density shown in magenta. **b**, varying thresholds of the substrate EM density, zoomed in on the length traversing the EA channel (box), with a modeled polyalanine sequence.

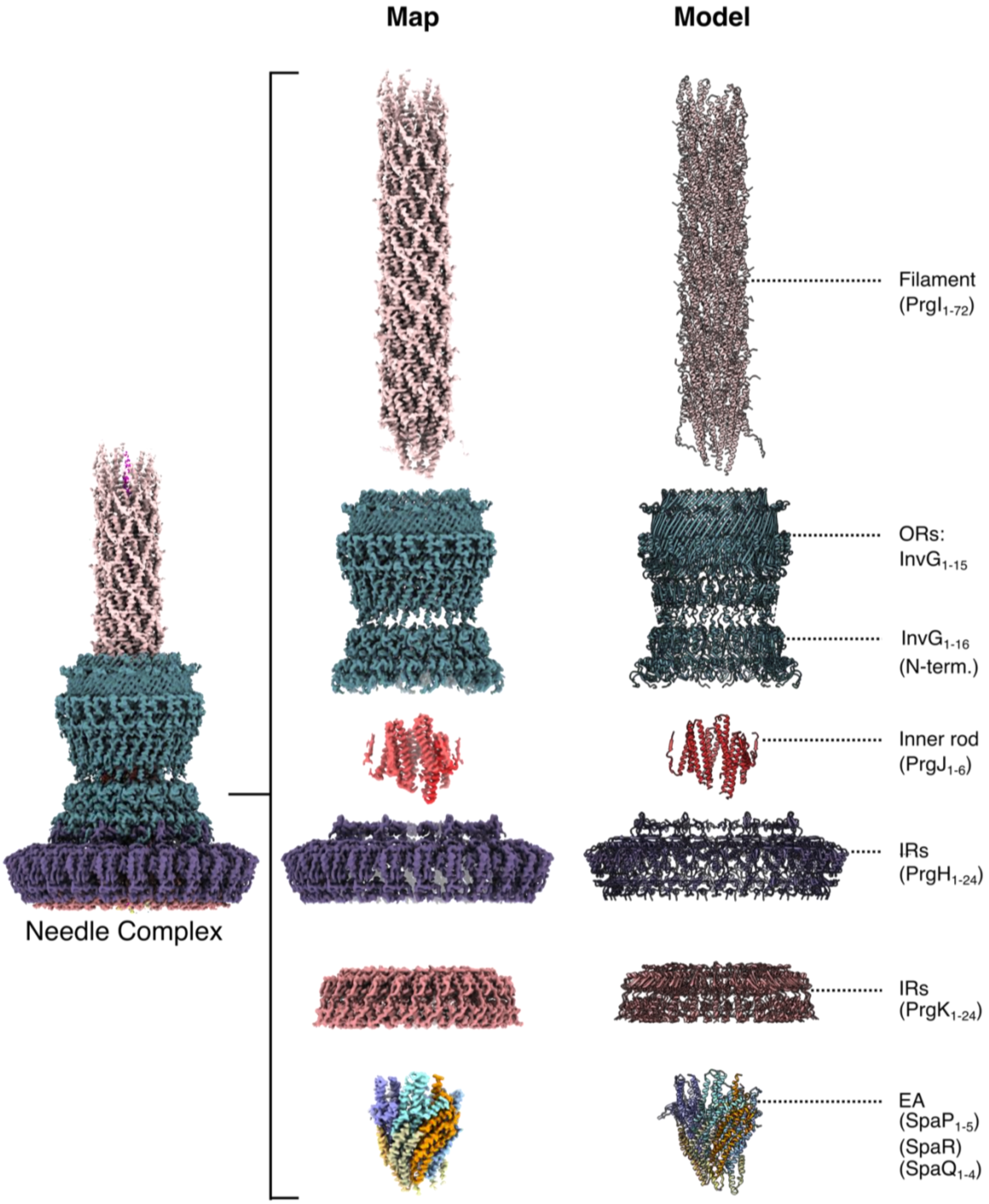


**Supplementary Figure 7: Needle complex assembly.** Extracted maps (left) and corresponding atomic models (right) of the export apparatus subcomplex containing SpaP, SpaQ and SpaR, the inner (IR; PrgK) and outer rings (OR; PrgH), the inner rod (PrgJ), the InvG ring with symmetry mismatch between the upper (C15) and lower (C16) OR component, and the filament (PrgI). Numbers correspond to the subunits present in each subcomplex.

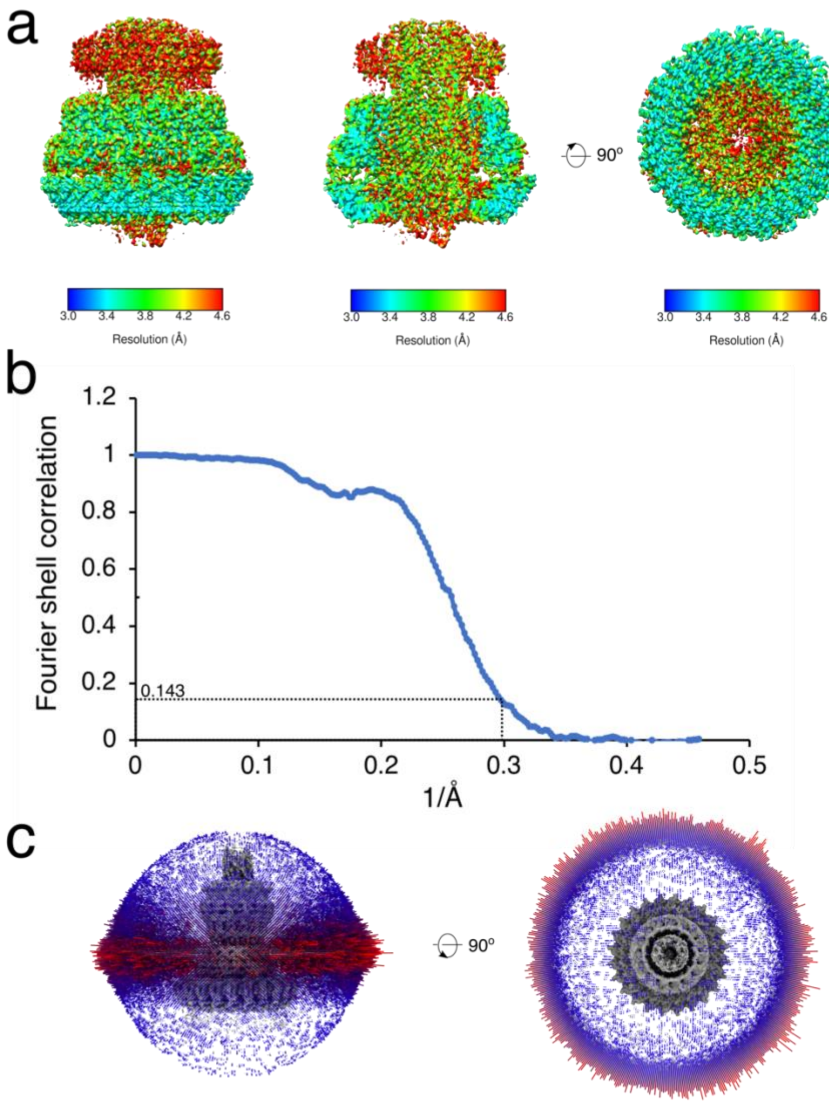


**Supplementary Figure 8: Single particle reconstruction of the apo-state needle complex from *S. enterica* sv. Typhimurium. a,** Local resolution estimation of the masked export apparatus and filament C1 map (left), a vertical cross section (middle) and a bottom view (right) calculated by ResMap[51]. **b,** Fourier Shell Correlation (FSC) of the masked C1 map. Global resolution at the gold-standard FSC threshold of 0.143 is ~3.32Å. **c**, Side (left) and top (right) view of the angular distribution plot of all needle complex particles that contributed to the final C1 map. Height and color (from blue to red) of the cylinder bars is proportional to the number of particles.

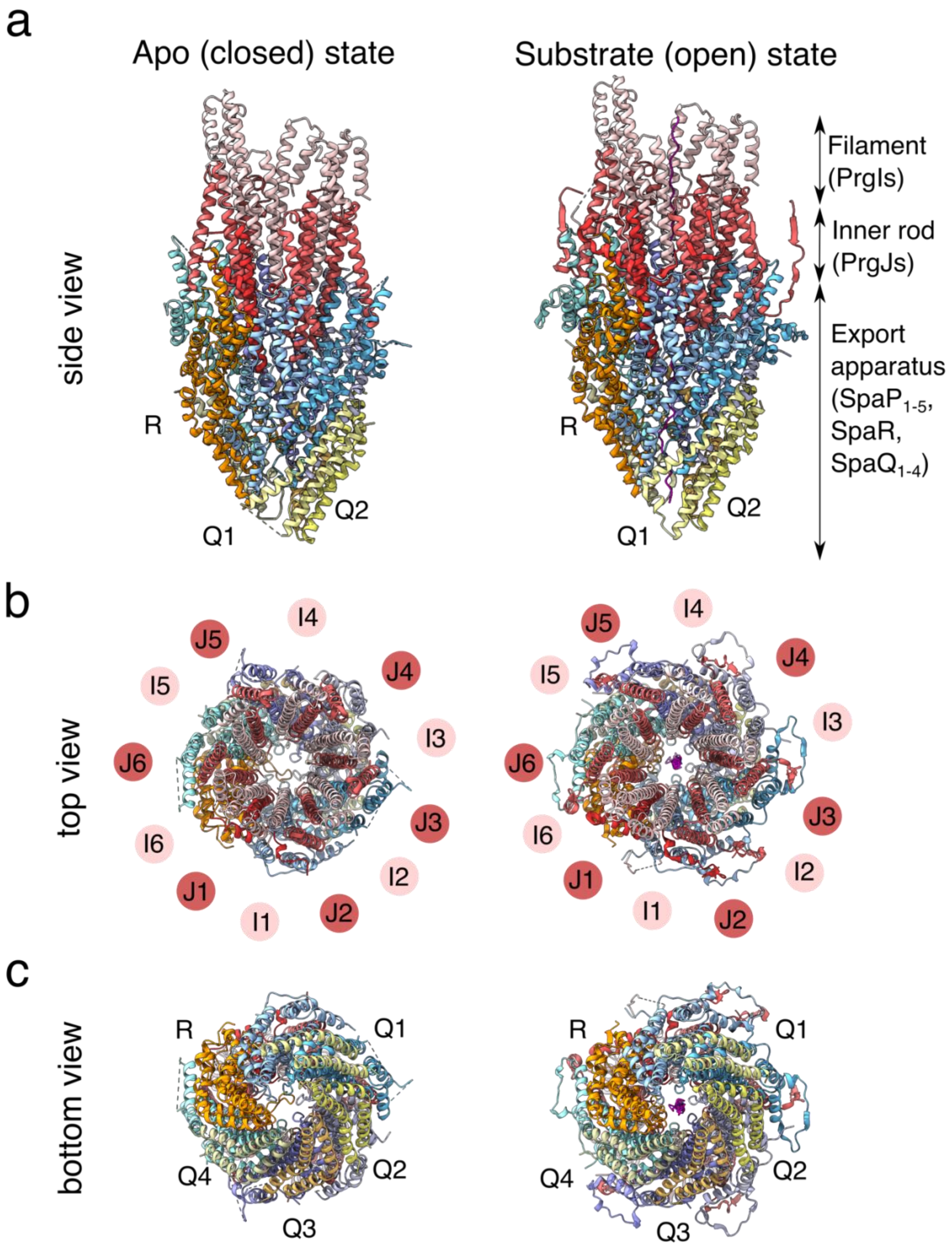


**Supplementary Figure 9: Comparison of apo (closed) and substrate-engaged (open) EA**

**complexes with assembled inner rod and filament base. a**, Side views. **b**, Top views. **c**, Bottom views. Color Code: SpaQ (Q1-Q4, yellow colors), SpaR (R, orange), SpaP (P1-P5, blue colors), PrgJ (J1-J6, red colors), PrgI (I1-I6, salmon), SptP3x-GFP substrate (magenta).

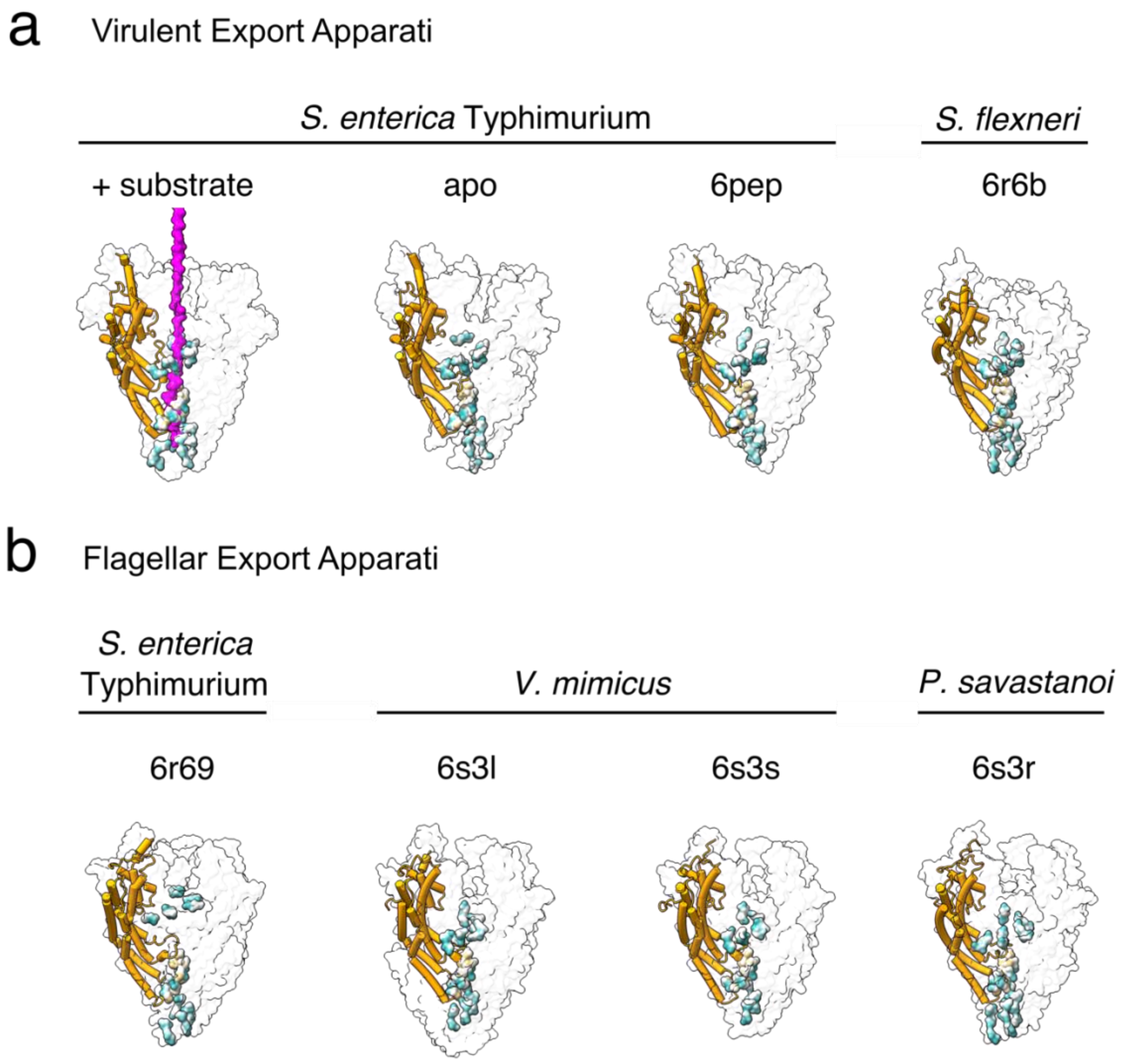


**Supplementary Figure 10: Conserved Q1-belt/M-gate/Q2-belt architecture in EAs from virulent and flagellar T3SSs. a**, Transparent surface representations of virulent EA proteins with SpaR homologues shown as cylinders colored in orange. Side chain surfaces of residues shaping the three EA interfaces, Q1-belt, M-gate and Q2-belt are highlighted where color represents hydrophobicity (green: hydrophilic; white: neutral; gold: hydrophobic). **b**, Representation style as in (a) but for published structures of flagellar EAs. SpaR homologue FliR colored in orange. Labels are PDB accession codes.

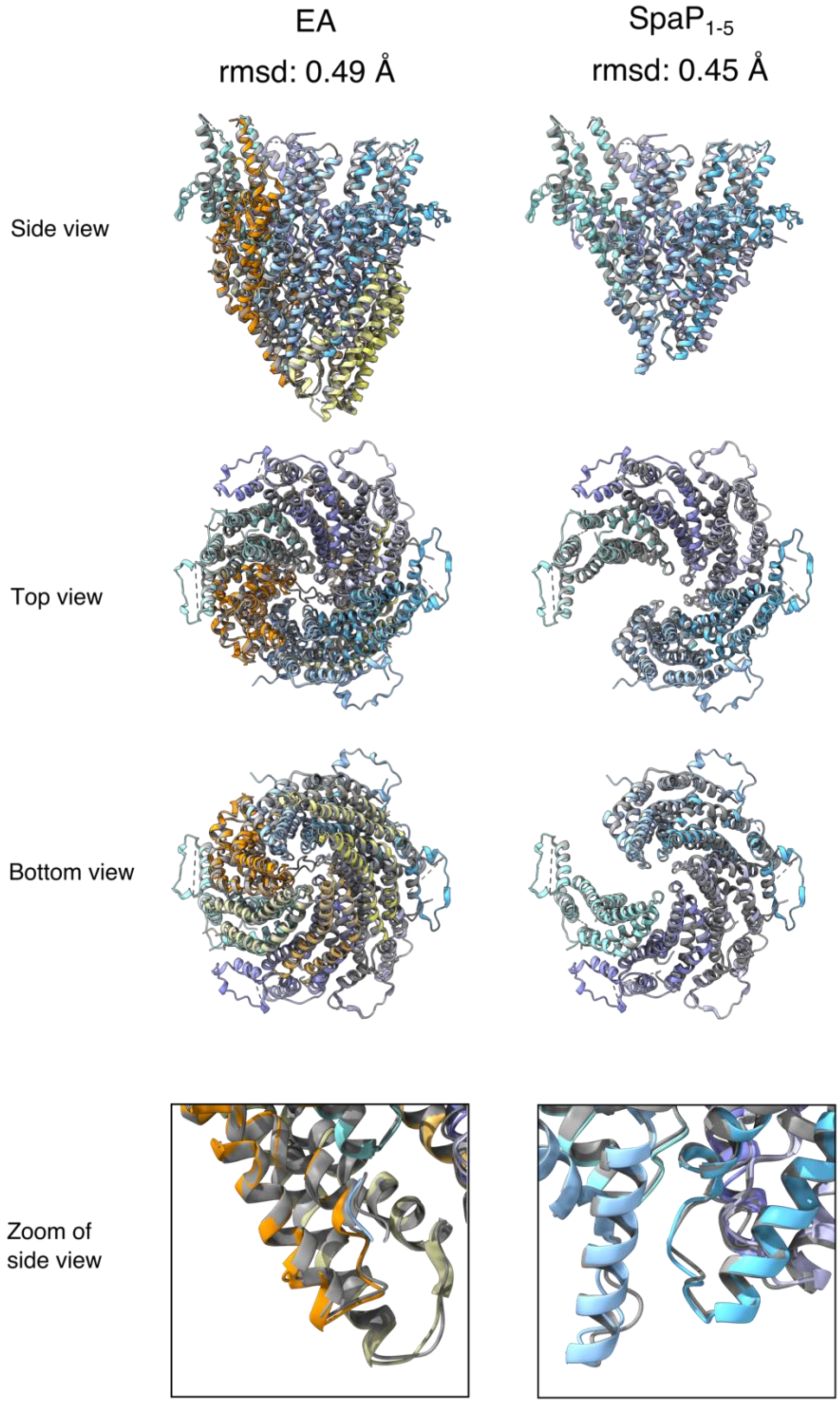

**Supplementary Figure 11: Superposition of closed (apo-state) and open (substrate-engaged) EA structures.** Proteins shown in ribbon diagrams with the apo-state EA colored in grey. Proteins of substrate-engaged EA are colored in orange (SpaR), blue colors ($SpaP_{1-5}$) and yellow colors ($SpaQ_{1-4}$). EA (SpaP:Q:R) complexes align with a root mean square deviation (rmsd) of 0.49Å (left). $SpaP_{1-5}$ align with a rmsd of 0.45Å (right).

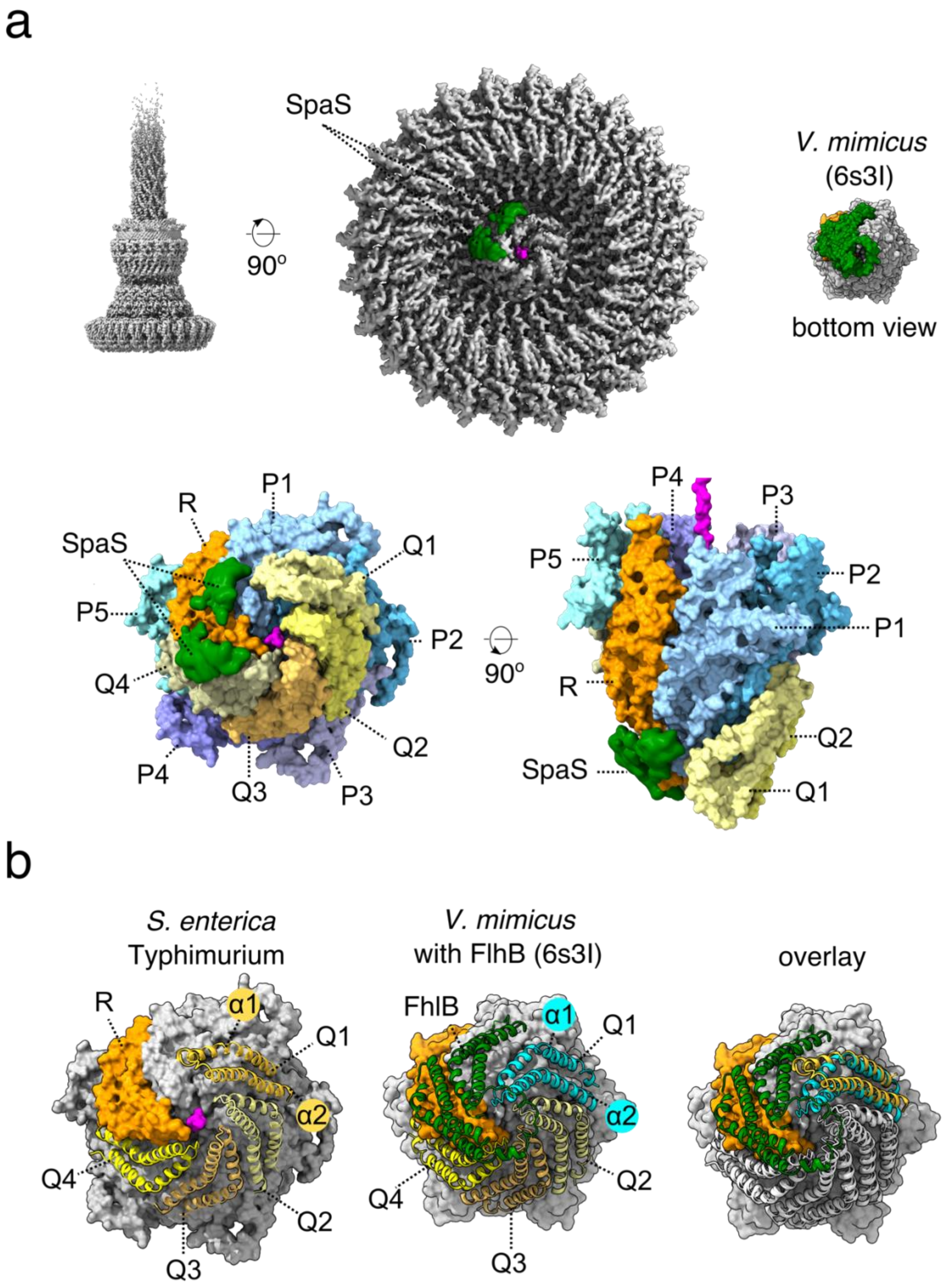


**Supplementary Figure 12: Q1-belt portal of the EA contains SpaS and has a displaced SpaQ$_1$ when compared to the flagellar homologue from *Vibrio mimicus*.** **a**, EM density (green)

corresponding to SpaS N-terminal helices seen in the substrate-engaged needle complex map at high threshold (middle) and the recombinant EA complex (6s3l) from *Vibrio mimicus* (right)[13]. Density was segmented from a smoothed map using Chimera Segger and placed on the map of the needle complex[52,53]. Below, SpaS density placed on the SptP3x-GFP-containing EA model shown in surface representation. Color code: SpaP (blue, P1-5), SpaR (orange), SpaQ (yellow, Q1-4). **b**, Left: Surface representation of EAs with SpaP proteins colored in grey and SpaR in orange. SpaQs$_{1-4}$ depicted as cartoon ribbons (yellow). Surface of the substrate model is displayed in magenta. α-helices of SpaQ$_1$ are labeled. Middle: Depiction of *V. mimicus* flagellar EA from Kuhlen *et al.*, 2020 (PDB: 6s3l) with surfaces of FliPs (SpaP homologue) and FliR (SpaR homologue) shown in grey and orange, respectively[13]. FliQs (SpaQ homologue) depicted as yellow cartoon ribbons. α-helices of FliQ$_1$ are labeled. Right: Superposition of SpaQs from substrate-engaged EA complex onto the *V. mimicus* EA with corresponding FliQ$_1$ highlighted in cyan. SpaQ$_{1-4}$ and FliQ$_{1-4}$ superimpose with a root mean square deviation (rmsd) of 1.27Å.

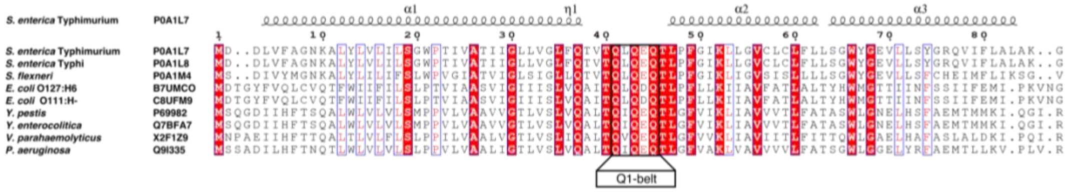


**Supplementary Figure 13: Multiple sequence alignment of SpaQ homologues.** *S. enterica* sv. Typhimurium (P0A1L7), *S. enterica* sv. Typhi (P0A1L8), *S. flexneri* (P0A1M4), EPEC (B7UMC0), EHEC (C8UFM9), *Y. pestis* (P69982), *Y. enterocolitica* (Q7BFA7), *V. parahaemolyticus* (X2F1Z9), *P. aeruginosa* (Q9I335). Uniprot accession numbers indicated. Residues involved in shaping the Q1-belt in SpaQ in *S. enterica* sv. Typhimurium (Gln41, Gln43 and Gln45) and other homologues are boxed out. Alignment performed using T-Coffee and displayed with EsPript 3.0 server[54,55].

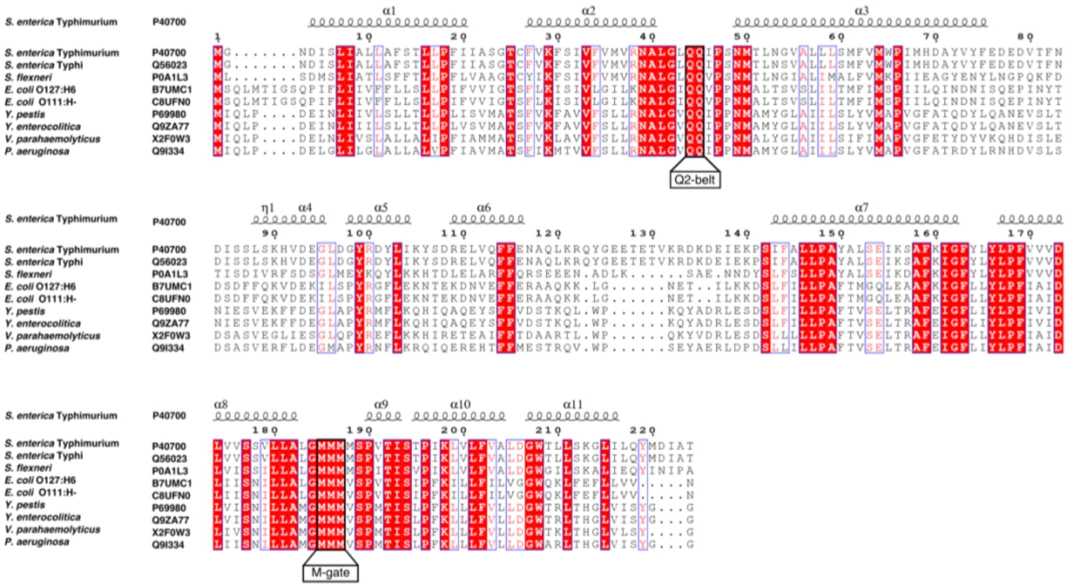


**Supplementary Figure 14: Multiple sequence alignment of SpaP homologues.** *S. enterica* sv. Typhimurium (P40700), *S. enterica* sv. Typhi (Q56023), *S. flexneri* (P0A1L3), EPEC (B7UMC1), EHEC (C8UFN0), *Y. pestis* (P69980), *Y. enterocolitica* (Q9ZA77), *V. parahaemolyticus* (X2F0W3), *P. aeruginosa* (Q9I334). Uniprot accession numbers are indicated. Residues involved in shaping the Q2-belt and M-gate in SpaP in *S. enterica* sv. Typhimurium (Gln44, Gln45 and Met185-187) and other homologues are boxed out. Alignment performed using T-Coffee, and displayed with EsPript 3.0 server[54,55].

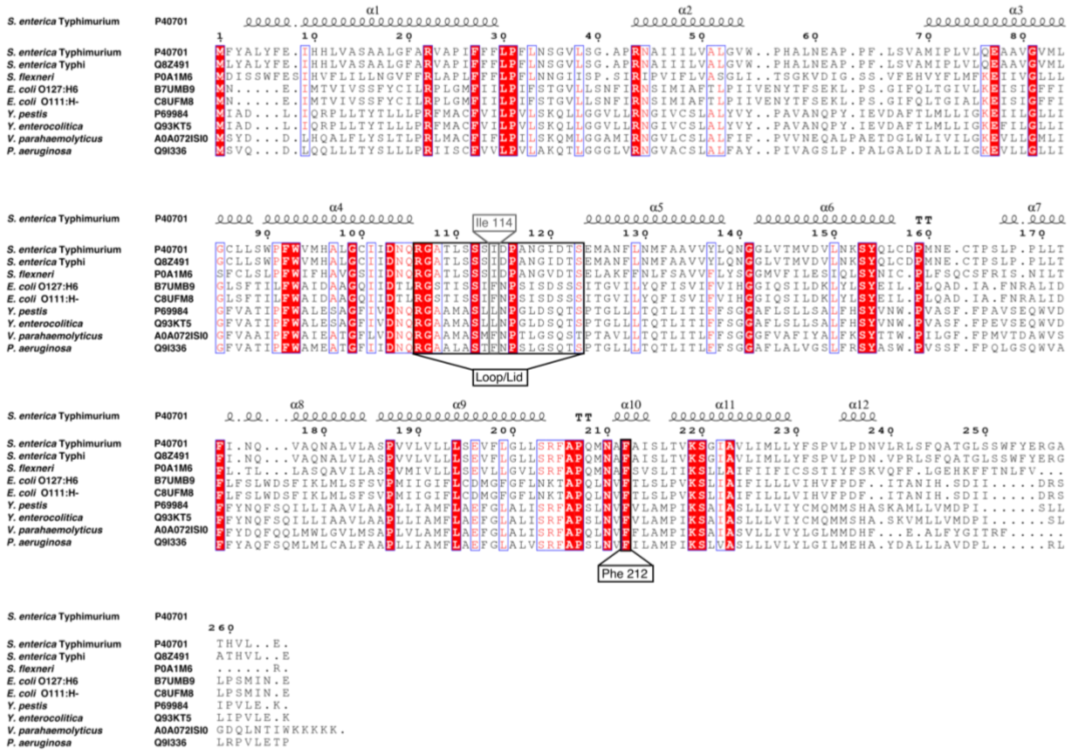


**Supplementary Figure 15: Multiple sequence alignment of SpaR homologues.** *S. enterica* sv. Typhimurium (P40701), *S. enterica* sv. Typhi (Q8Z491), *S. flexneri* (P0A1M6), EPEC (B7UMB9), EHEC (C8UFM8), *Y. pestis* (P69984), *Y. enterocolitica* (Q93KT5), *V. parahaemolyticus* (A0A072ISI0), *P. aeruginosa* (Q9I336). Uniprot accession numbers are indicated. Residues involved in shaping the loop/lid, Ile114, and Phe212 of SpaR in *S. enterica* sv. Typhimurium and other homologues, are boxed out. Alignment performed using T-Coffee and displayed with EsPript 3.0 server[54,55].

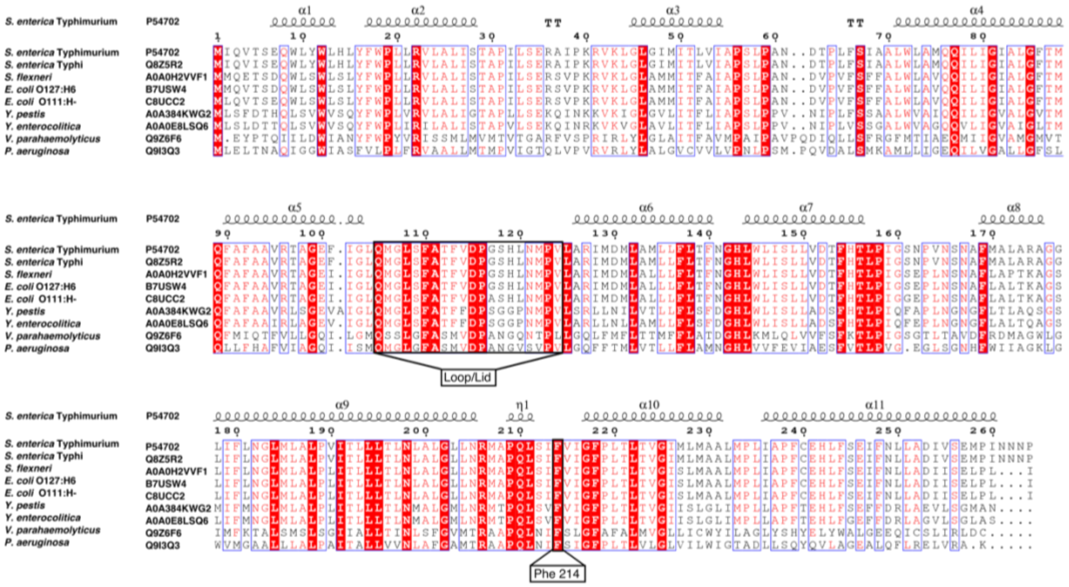


**Supplementary Figure 16: Multiple sequence alignment of FliR homologues.** *S. enterica* sv. Typhimurium (P54702), *S. enterica* sv. Typhi (Q8Z5R2), *S. flexneri* (A0A0H2VVF1), EPEC (B7USW4), EHEC (C8UCC2), *Y. pestis* (A0A384KWG2), *Y. enterocolitica* (A0A0E8LSQ6), *V. parahaemolyticus* (Q9Z6F6), *P. aeruginosa* (Q9I3Q3). Uniprot accession numbers are indicated. Residues involved in shaping the FliR loop/lid and Phe214 are boxed out and correspond to the homologous loop/lid and Phe212 of SpaR in *S. enterica* sv. Typhimurium. Alignment performed using T-Coffee and displayed with EsPript 3.0 server[54,55].

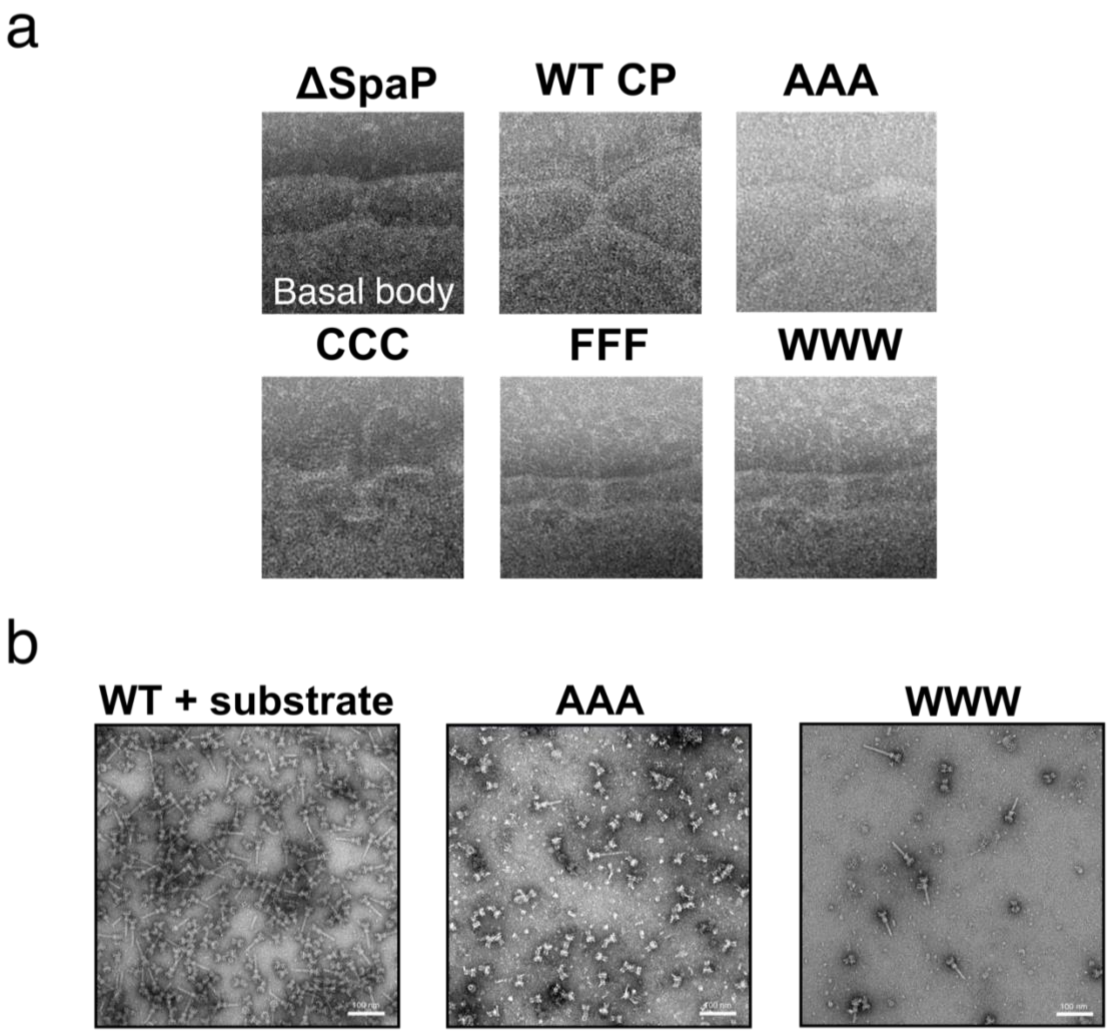


**Supplementary Figure 17: Visualization of needle complexes in *Salmonella* SpaP M-gate mutants.** **a**, Representative micrographs of basal bodies (ΔSpaP) and needle complexes (AAA, CCC, FFF, WWW) in osmotically-shocked cells from SpaP knockout strains complemented with the corresponding SpaP M-gate mutations. **b**, Micrographs of needle complex purifications from *Salmonella* expressing the substrate SptP3x-GFP (left) or $SpaP^{KO}$ knockout strain complemented with $SpaP^{AAA}$ (middle) or $SpaP^{WWW}$ (right). Samples were imaged via negative stain TEM using 2% w/v PTA. Needle-less basal bodies correspond to incomplete assembly states of needle complexes.

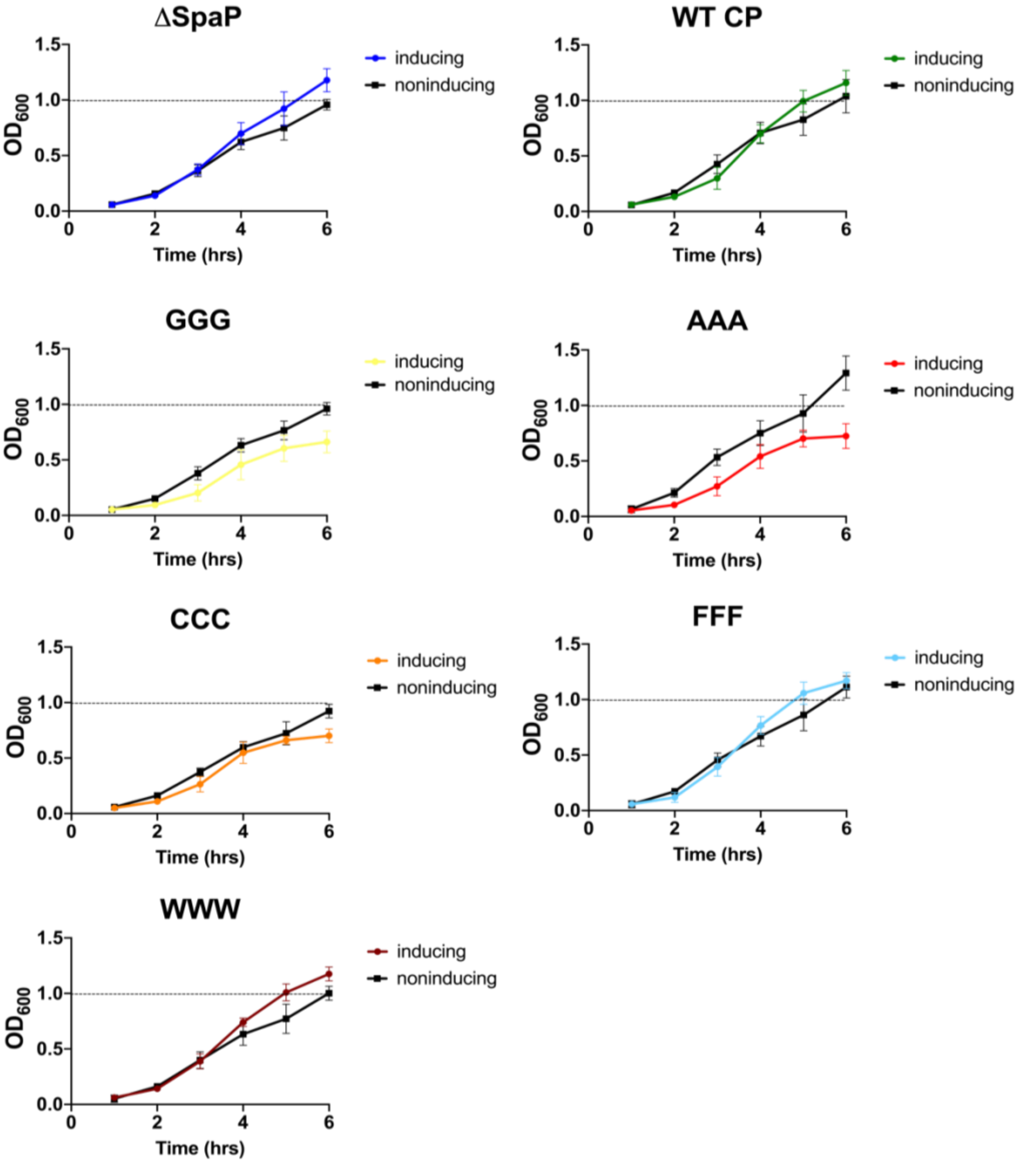


**Supplementary Figure 18: Growth curves of *Salmonella* SpaP knockout strains complemented with SpaP harboring M-gate mutations.** Each chart plots $OD_{600}$ of a 6 hour time course for each strain. All strains are SpaP knockout and WT CP is complemented with WT SpaP. Black and colored lines represent growth under non-inducing and inducing conditions, respectively. Plotted values are the average from 9 independent measurements. Error bars represent SD.

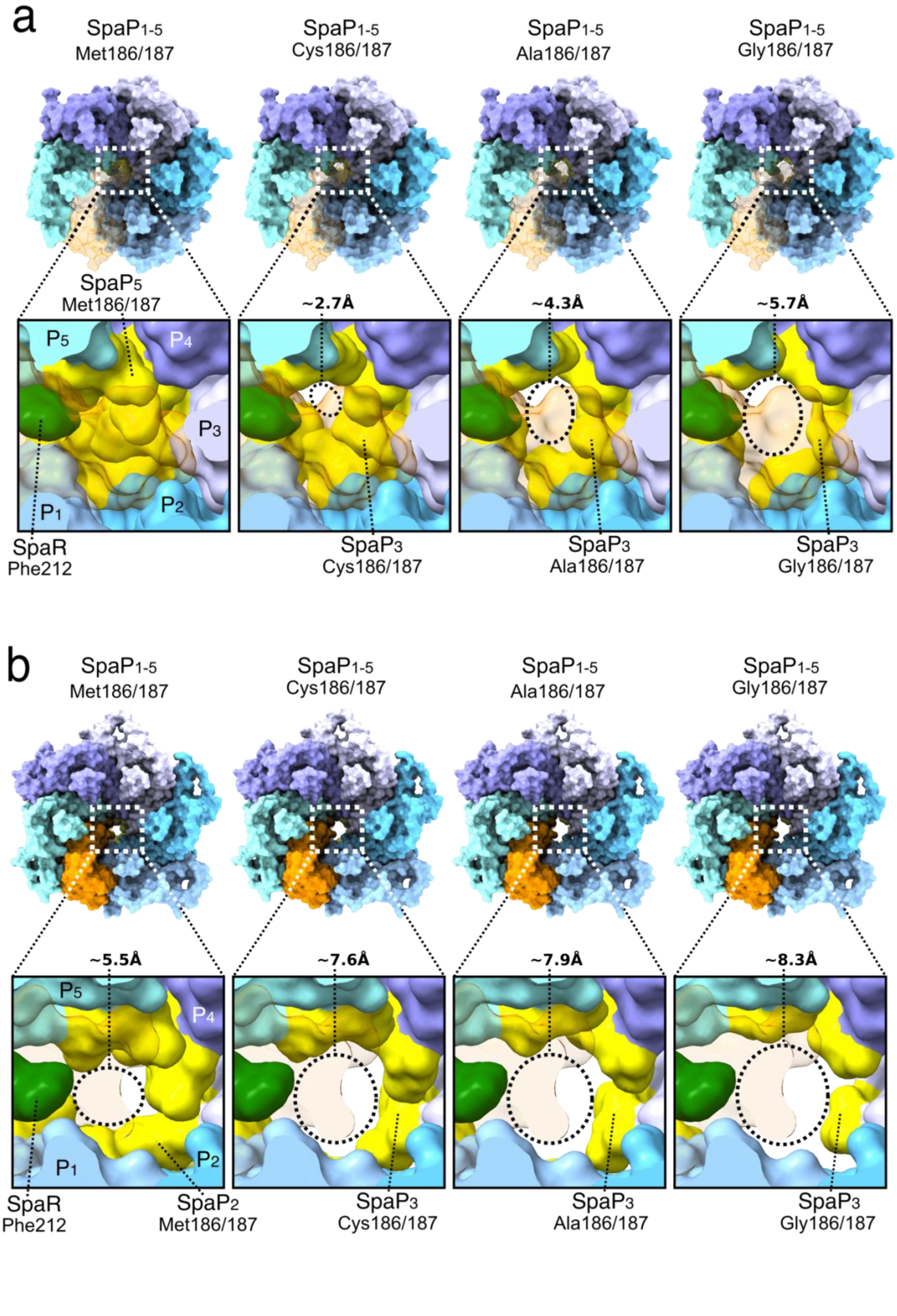
a
SpaP1-5
Met186/187
SpaP1-5
Cys186/187
SpaP1-5
Ala186/187
SpaP1-5
Gly186/187
SpaP5
Met186/187
~2.7Å
~4.3Å
~5.7Å
P5
P4
P3
P1
P2
SpaR
Phe212
SpaP3
Cys186/187
SpaP3
Ala186/187
SpaP3
Gly186/187
b
SpaP1-5
Met186/187
SpaP1-5
Cys186/187
SpaP1-5
Ala186/187
SpaP1-5
Gly186/187
~5.5Å
~7.6Å
~7.9Å
~8.3Å
P5
P4
P1
P2
SpaR
Phe212
SpaP2
Met186/187
SpaP3
Cys186/187
SpaP3
Ala186/187
SpaP3
Gly186/187

**Supplementary Figure 19: Mutations of methionines 186 and 187 to cysteines, alanines or glycines increase the pore size of the M-gate. a**, Top views onto the apo-state EA complex with $SpaP_{1-5}$ Met186 and Met187 mutated to either cysteine, alanine or glycine. Approximate diameters of the M-gates were calculated from modeled surfaces. Surface of SpaR (with the exception of SpaR Phe212 shown in green) is transparent to aid visualization of the M-gate. Residues 186 and 187 are colored in yellow. **b**, Top views onto the substrate-state EA complex with $SpaP_{1-5}$ mutated to either cysteine, alanine or glycine. Approximate diameters of the M-gates were calculated from modeled surfaces. The substrate has been removed to aid visualization.

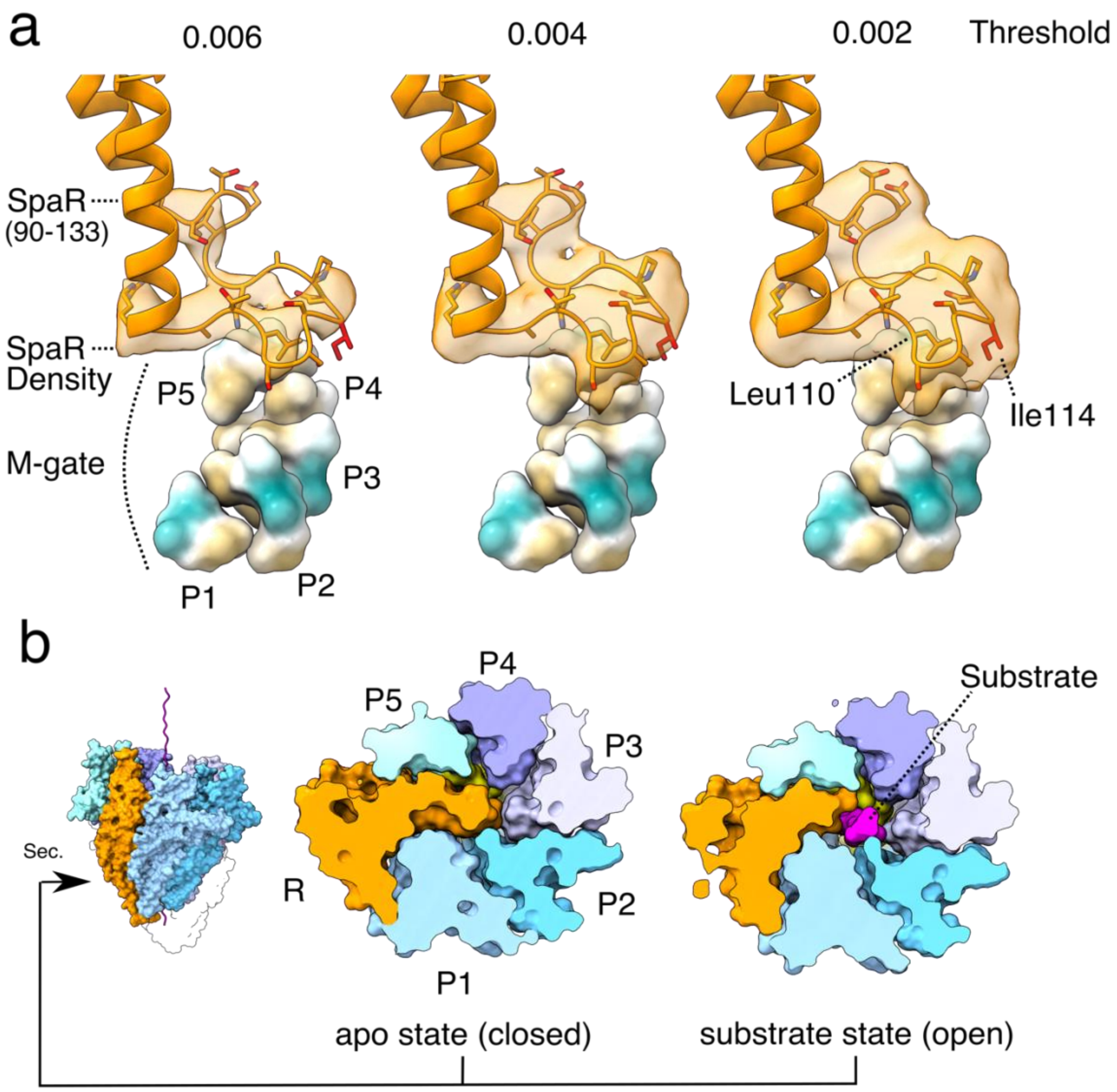


**Supplementary Figure 20: A unique loop of residues 106-123 of SpaR seals the EA translocation channel. a,** The closed SpaR loop/lid is shown with a filtered density (B-factor of 100) and modeled residues. The SpaP M-gate methionine residues are shown below as surface representations and colored by hydrophobicity. **b,** Left: overview of the EA with the horizontal position marked corresponding to cross sections of the SpaR loop through the apo-state EA complex (middle) and substrate-engaged EA complex (right). Proteins are depicted as surface representations. SpaP$_{1\text{-}4}$ is depicted in blue, SpaR in orange, and SpaQ1-4 is transparent. The substrate is depicted in magenta and the M-gate methionines are colored yellow.

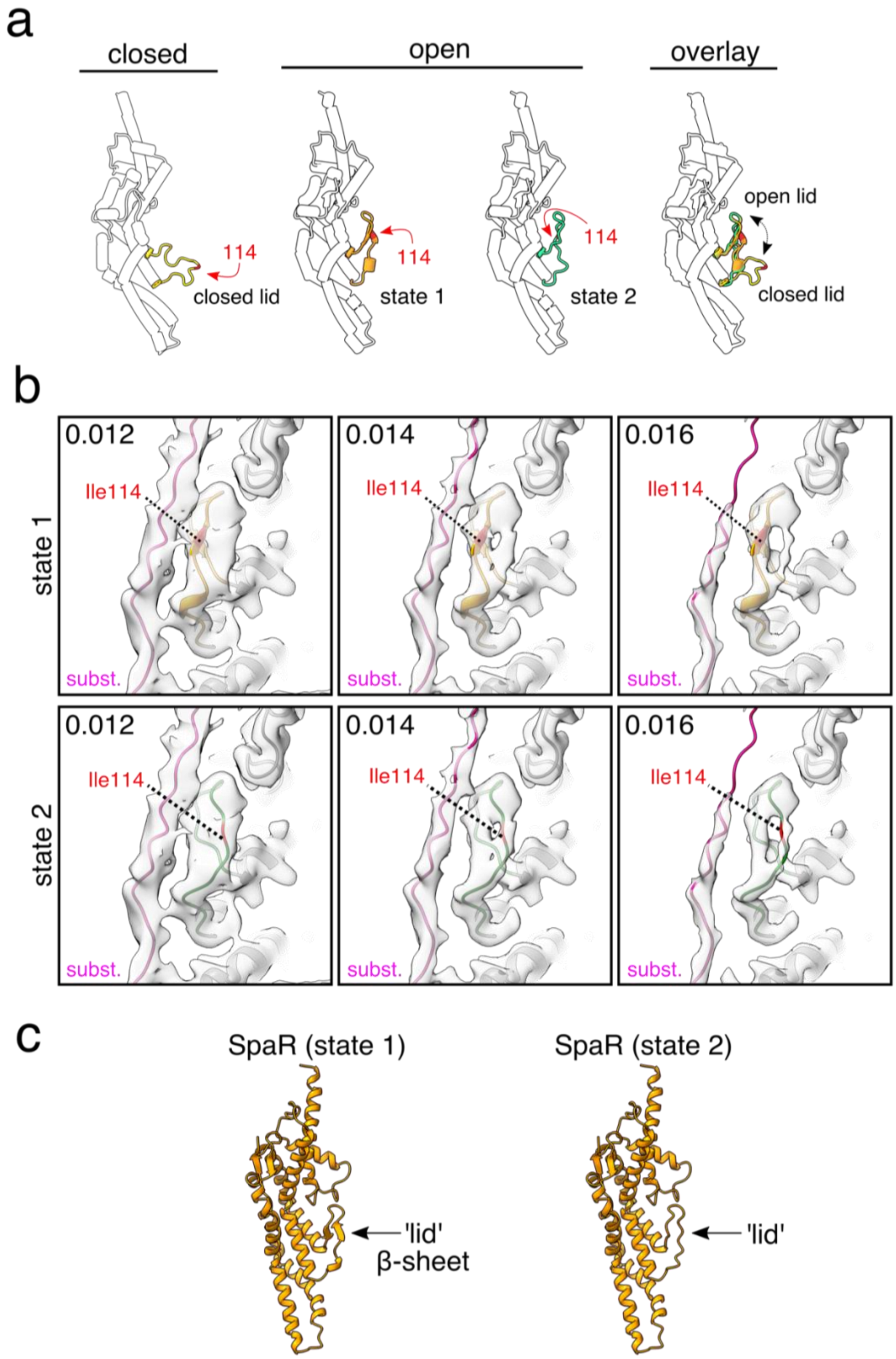
a
closed
open
overlay
114
closed lid
114
state 1
114
state 2
open lid
closed lid
b
state 1
state 2
0.012
0.014
0.016
Ile114
subst.
c
SpaR (state 1)
SpaR (state 2)
'lid'
β-sheet
'lid'

**Supplementary Figure 21: SpaR loop 106-123 forms two distinct conformations. a**, Transparent cylinder representation of SpaR with its lid in closed (left; yellow) or two distinct open conformations (middle; state 1 is colored orange, state 2 is green). Overlay between all three conformations (right) illustrates the movement of the loop from a horizontal (closed state) to vertical positions (open states). **b**, Panels show EM densities with increasing map threshold levels (numbers). Model is depicted in ribbon diagram representation with the substrate (subst.) shown in magenta and SpaR loop conformational states one and two in orange or green, respectively. Position of Ile114 is highlighted in red. **c**, Ribbon diagrams of SpaR with loop residues 106-123 in the two different conformations where in state 1, the SpaR loop is stabilized by a short antiparallel β-sheet.

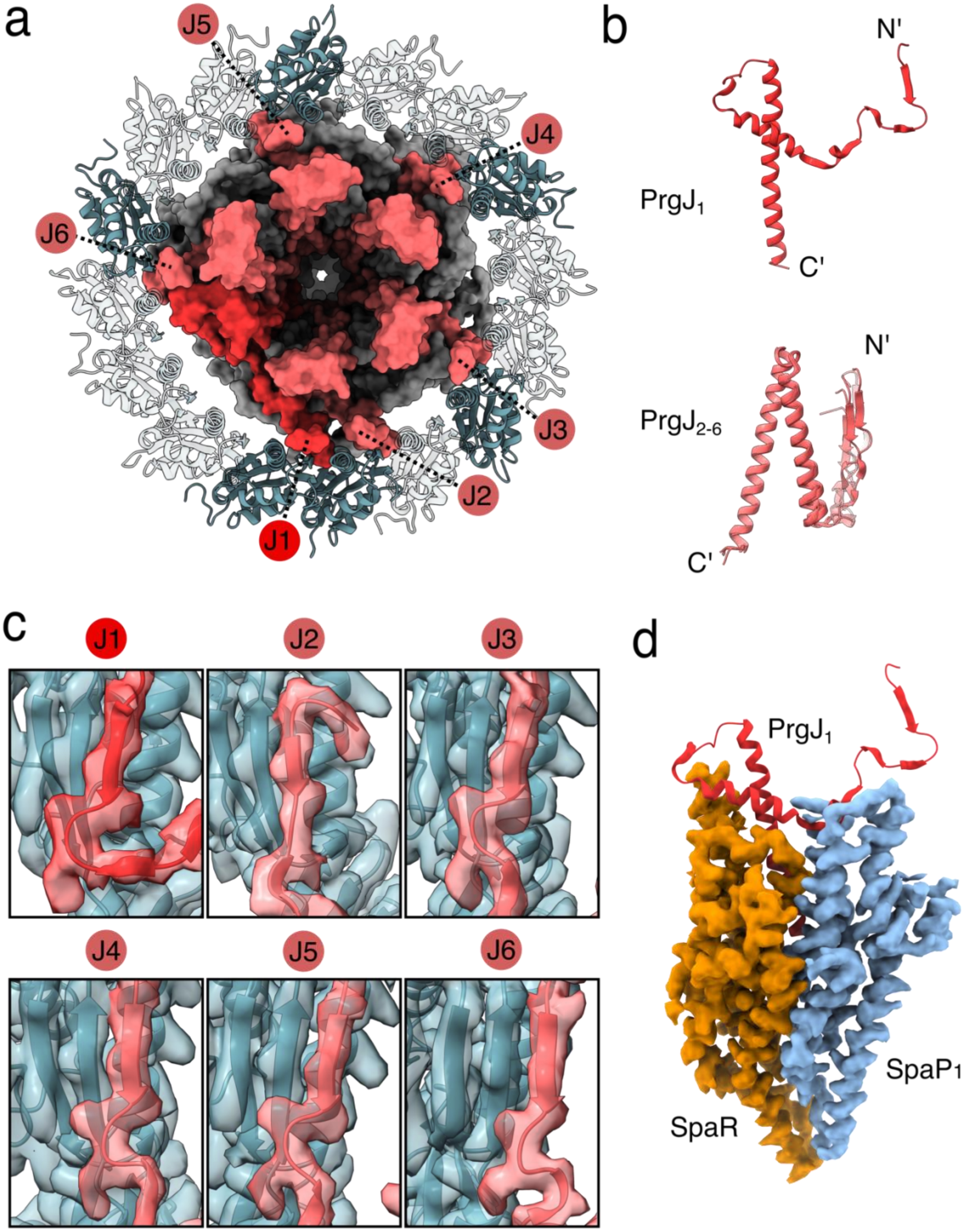
a
J5
J4
J6
J3
J2
J1
b
N'
PrgJ1
C'
N'
PrgJ2-6
C'
c
J1
J2
J3
J4
J5
J6
d
PrgJ1
SpaP1
SpaR

**Supplementary Figure 22: PrgJ proteins contact EA and InvG proteins to stabilize the inner rod in the confinement of the basal body. a**, Top view of the EA and inner rod together in surface representation and with the lower part of the InvG 16-mer ring in ribbon diagram representation (also see Supplementary Fig. 7). InvG monomers forming contacts with $PrgJ_{1-6}$ N-termini are colored dark green. **b,** Comparison of the unique fold of $PrgJ_1$ (top) and superimposed $PrgJ_{2-6}$ (bottom) shown as ribbon diagrams. **c,** Close up views of $PrgJ_{1-6}$ N-termini forming β-sheet complementation with subunits of the InvG 16-mer ring, as shown in (**a**), depicted as ribbon diagrams with the corresponding EM density. **d,** The unique fold of $PrgJ_1$ shown as a red ribbon diagram, interfacing SpaR shown as EM density in orange and traversing $SpaP_1$ also shown as EM density in blue.

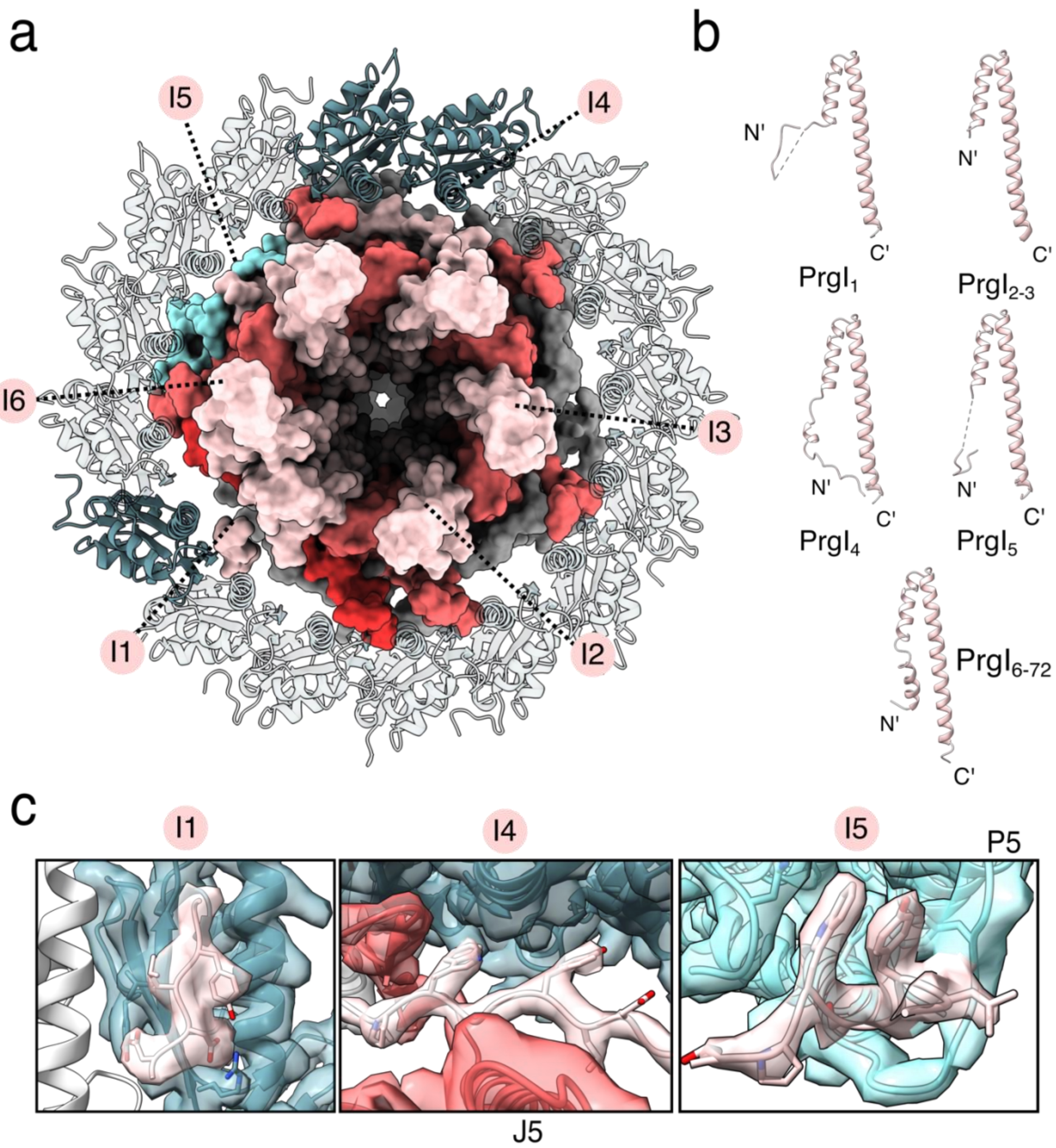


**Supplementary Figure 23: PrgI proteins polymerize on top of the inner rod and contact EA and InvG proteins to stabilize the filament in the confinement of the basal body. a**, Top view onto the EA with molecular surfaces of $SpaP_5$ in blue and $SpaP_{1\text{-}4}$ in grey, PrgJ in red and PrgI in salmon. InvG N-terminal domains are depicted as ribbon diagrams. InvG monomers forming contacts with $PrgI_{1,4,5}$ are colored in dark green **b**, Ribbon diagrams of PrgI proteins illustrating structural differences between $PrgI_1$, $PrgI_{2-3}$, $PrgI_4$, $PrgI_5$ and $PrgI_{6-72}$. **c**, Interfaces of $PrgI_{1,4-5}$ N-termini with InvG ($PrgI_1$, $PrgI_4$), $PrgJ_5$ ($PrgI_4$) and $SpaP_5$ ($PrgI_5$). Surfaces represent EM density.

| **Sample** | | **Needle complex apo state** | **Needle complex substrate-trapped** |
|---|---|---|---|
| | Buffer | 10 mM Tris-HCl, pH 8.0, 0.5 M NaCl, 5 mM EDTA, 0.1% w/v LDAO | |
| | Vitrification | Thermo Fisher Scientific Vitrobot mark III | |
| | Grid | Quantifoil with floated carbon | |
| | EMBD ID | 11780 | 11781 |
| **Data Collection** | Magnification, nominal | 130kx | 81kx |
| | Energy filter slit, (eV) | 15 | 10 |
| | Defocus range set for data collection (µm) | 0.3-5.2 | 0.3-5.2 |
| | Voltage (kV) | 300 | 300 |
| | Microscope | Thermo Fisher Scientific Titan Krios | Thermo Fisher ScientificTitan Krios |
| | Camera | Gatan K2 | Gatan K3 |
| | Frame exposure time (sec) | 0.2 | 0.06 |
| | Dose per frame (e-/Å2) | 1.26 | 1.06 |
| | Number of movie frames | 25 | 50 |
| | Pixel size (Å) | 1.09 | 1.1 (0.55 super resolution) |
| | Total electron dose, (e-/Å2) | 31.5 | 53 |
| | Number of micrographs collected | 10433 | 14450 |
| **Data Processing** | Number of picked coordinates (crYOLO) | 171130 + 62972 (Graphene Oxide) | 837325 |

| | | |
|---|---|---|
| Number of particles for polishing | 86473 | 296521 |
| Particle number in final map | 54491 | 77411 |
| Resolution with masking (Å) | 3.36 | 3.32 |
| Applied symmetry | C1 | C1 |
| Calculated resolution range with ResMap (Å) | 2.4-4.5 | 2.4-4.5 |
| B-factor (manual) | -30 | -30 |

| | | | **SpaR state 1** | **SpaR state 2** |
|---|---|---|---|---|
| **Model Building** | PDB ID | 7agx | 7ah9 | 7ahi |
| | Clashscore, all atoms | 2.64 (98th percentile) | 2.36 (99th percentile) | 2.35 (99th percentile) |
| | Poor rotamers | 2 – 0.08% | 1 – 0.00% | 1 – 0.00% |
| | Favoured rotamers | 2579 – 98.77% | 21192 – 98.38% | 21195 – 98.40% |
| | Ramachandran outliers | 0 – 0.00% | 0 – 0.00% | 0 – 0.00% |
| | Ramachandran favored | 2902 – 98.94% | 24510 – 98.19% | 24508 – 98.19% |
| | Rama distribution Z-score | -2.48 ± 0.13 | -1.75 ± 0.05 | -1.75 ± 0.05 |
| | MolProbity score | 1.05 (100th percentile) | 1.02 (100th percentile) | 1.01 (100th percentile) |
| | Cβ deviations > 0.25Å | 0 – 0.00% | 0 – 0.00% | 0 – 0.00% |
| | Bad bonds: | 0/24089 – 0.00% | 0/203687 – 0.00% | 2/203687 – 0.00% |
| | Bad angles: | 0/32712 – 0.00% | 21/276188 – 0.01% | 3/276188 – 0.00% |
| | Cis Prolines: | 7/96 – 7.29% | 44/1044 – 4.21% | 44/1044 – 4.21% |
| | CaBLAM outliers | 12 – 0.40% | 220 – 0.9% | 220 – 0.9% |
| | CA Geometry outliers | 4 – 0.14% | 86 – 0.35% | 85 – 0.35% |

| | | | |
|---|---|---|---|
| Chiral volume outliers | 0/3924 | 0/31002 | 0/31002 |
| EMRinger score | 3.55 | 3.36 | 3.37 |

**Supplementary Table 1.** Cryo-EM data collection, processing and model building summary.

| Structure | Conformational state 1 | Conformational state 2 |
|---|---|---|
| interface area | 679Å | 694Å2 |
| ΔG: | -4.4 kcal/mol | -4.1 kcal/mol |
| ΔG p-value | 0.86 | 0.85 |
| Complexation Significance score | 0 | 0 |

**Supplementary Table 2. PDBePISA analysis of SpaR loop/lid conformational states 1 and 2[56].**
